# Combined social and spatial coding in a descending projection from the prefrontal cortex

**DOI:** 10.1101/155929

**Authors:** M. Murugan, M. Park, J. Taliaferro, H.J. Jang, J. Cox, N.F. Parker, V. Bhave, A. Nectow, J.W. Pillow, I.B. Witten

## Abstract

Social interactions are crucial to the survival and well-being of all mammals, including humans. Although the prelimbic cortex (PL, part of medial prefrontal cortex) has been implicated in social behavior, it is not clear which neurons are relevant, nor how they contribute. We found that the PL contains anatomically and molecularly distinct subpopulations of neurons that target 3 downstream regions that have been implicated in social behavior: the nucleus accumbens (NAc), the amygdala, and the ventral tegmental area. Activation of NAc-projecting PL neurons (PL-NAc), but not the other subpopulations, decreased preference for a social target, suggesting an unique contribution of this population to social behavior. To determine what information PL-NAc neurons convey, we recorded selectively from them, and found that individual neurons were active during social investigation, but only in specific spatial locations. Spatially-specific inhibition of these neurons prevented the formation of a social-spatial association at the inhibited location. In contrast, spatially nonspecific inhibition did not affect social behavior. Thus, the unexpected combination of social and spatial information within the PL-NAc population appears to support socially motivated behavior by enabling the formation of social-spatial associations.

## Introduction

Pioneering research on social behavior in animal models has focused on identifying neural circuitry underlying aggression [1–12] and sexual behavior [13–20]. In contrast, relatively little is known about neural substrates of non-aggressive same-sex social interactions, despite the fact that such behaviors are central to most mammalian species, including humans, and are of relevance to a number of neuropsychiatric diseases [21–23].

A common non-aggressive and non-sexual social behavior in mice and other rodents is the tendency to investigate novel same-sex conspecifics, a behavior characterized by sniffing and grooming. This investigation behavior is typically rewarding, as evidenced by the fact that mice are likely to return to the location in which they previously encountered a conspecific [24,25], and the fact that the behavior generates release of dopamine in the nucleus accumbens (NAc) [26–29].

Despite the robust and rewarding nature of non-aggressive social investigation, understanding its neural basis has been challenging, in part due to the complexity of the underlying circuitry. In mice, a large number of distributed but interconnected brain areas have been implicated [2,5,30–32], including the prelimbic cortex (PL, part of the medial prefrontal cortex) [33–44] and several of its projection targets (e.g. the amygdala, ventral tegmental area, NAc) [26,42,45–48]

In particular, activation of the PL causes mice to spend significantly less time investigating a conspecific, suggesting an important role of this region in social behavior [33]. However, inhibition of the endogenous activity in the same region results in no observable change in behavior [33]. This discrepancy raises the possibility that experimental activation of PL reduces social investigation by producing unnatural patterns of activation of projection targets, rather than by replicating activity that occurs endogenously during the behavior [49].

To address this possibility, and more broadly to clarify the nature of PL’s contribution to social behavior, we manipulated and recorded from projection-defined subpopulations of PL neurons. We first sought to determine which descending projection from PL impacts social investigation when activated. We found that activation of PL neurons that project to the NAc decreased social investigation, while other descending projections from PL did not have this effect. We then investigated if and how the endogenous activity within the PL-NAc subpopulation is modulated by social investigation. Cellular resolution imaging revealed that many neurons were active during social investigation, but surprisingly, this activity depended on the location of the social encounter. Based on this finding, we inhibited the activity in PL-NAc neurons in a spatially-specific manner. This experiment was motivated by the hypothesis that the combined social-spatial code in these neurons could guide social behavior by governing the formation of social-spatial associations, perhaps through synaptic plasticity in the NAc. Consistent with this hypothesis, spatially-specific inhibition of these neurons during social investigation prevented the formation of a preference for the location of a social encounter, while spatially non-specific inhibition had no effect on social investigation. Thus, PL-NAc neurons encode a combination of social and spatial information which appears to support learning to associate a spatial location with a social encounter.

## Results

### Anatomically and molecularly distinct neurons project from PL to the NAc, VTA, and amygdala

To address the nature of PL’s control over social behavior, we first asked if there are distinct subpopulations of neurons within PL that target major downstream regions [50-52] that have themselves been implicated in social behavior [26,45,46]. To this end, retrograde tracers were injected into either the nucleus accumbens (NAc), amygdala (Amyg) and ventral tegmental area (VTA) to enable visualization of PL neurons that project to each target region (“PL-NAc”, “PL-Amyg”, and “PL-VTA” subpopulations; Fig 1A-C; Supplementary Fig. 1A-C, n = Amyg: 390 neurons from 5 animals, NAc: n = 867 neurons, from 6 animals, VTA: n = 803 neurons from 7 animals). We observed a similar density of PL-NAc, PL-Amyg and PL-VTA neurons within the PL (Fig. 1D; Mean± S.E.M: NAc: 134±40, Amyg: 80±17 and VTA: 105±20 neurons/mm^2^; p=0.43 for 1-way ANOVA of cell density between the subpopulations). However, the 3 subpopulations occupied different cell layers, with the PL-NAc neurons positioned more medially than the PL-VTA neurons, and the PL-Amyg neurons distributed more broadly than the other two subpopulations (Fig. 1C,E, K-S tests to compare M/L distribution of the 3 populations in a pairwise manner, p< 0.0001 for all 3 comparisons, Bonferroni-Holm corrected; M/L cell distributions at 3 A/P locations in Supp. Fig. 1D-L). Consistent with the distinct laminar distributions, there was little overlap at the single cell level between the 3 subpopulations, as assessed by co-labelling of multiple retrograde tracers, indicating that PL-NAc, PL-Amyg and PL-VTA neurons are anatomically distinct subpopulations (Fig. 1F; Neuron overlap - NAc-Amyg: 6.09% or 19/312 from 2 animals, NAc-VTA: 3.65% or 35/959 from 3 animals, Amyg-VTA: 0.21% or 1/479 from 2 animals).

**Fig. 1.**
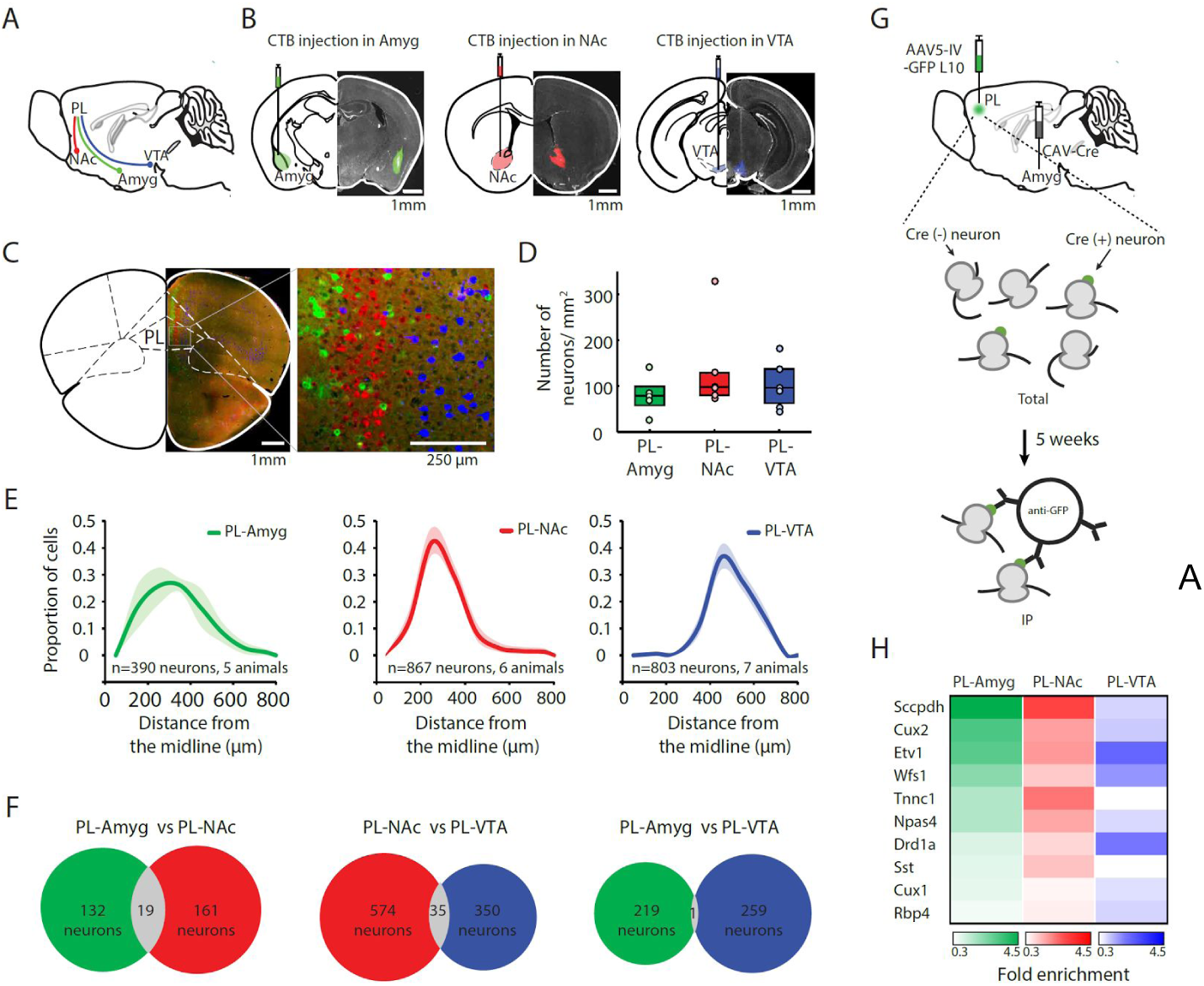
Anatomically and molecularly distinct subpopulations of PL neurons project to the NAc, VTA and Amygdala. **A.** A schematic of the mouse brain with subcortical projections of the PL to the Amyg (green), NAc (red), and the VTA (blue). **B.** Each brain section includes schematic (left) and histology (right) of CTB targeting to the Amyg (left panel), the NAc (center panel) and the VTA (right panel). Scale bar, 1 mm. **C.** coronal section of retrogradely labeled neurons in PL from CTB injections in Amyg (green), NAc (red) and VTA (blue). 2.2 m anterior bregma, scale bar 1 mm in main panel, 250 μm in inset. **D.** A similar number of PL neurons labeled with CTB from Amyg, NAc and VTA injections. (Mean ±S.E.M: 134±40 neurons/mm2 for NAc; 80±17 for Amyg; and 105±20 for VTA; p=0.43 for 1-way ANOVA comparing cell density between the subpopulations). In the box plots, the center indicates the median, and bottom and top edges indicate 25th and 75th percentiles, respectively. **E.** Quantification of the M/L distribution of PL-Amyg (green), PL-NAc (red) and PL-VTA (blue) neurons. Shading denotes S.E.M. The 3 subpopulations are distributed differently (K-S tests to compare M/L distribution of the 3 populations in a pairwise manner, p< 0.0001 for all 3 comparisons, Bonferroni-Holm corrected). **F.** Little overlap between the 3 projection populations at the individual cell level (Neuron overlap - NAc-Amyg: 6.09% or 19/312 neurons from 2 animals, NAc-VTA: 3.65% or 35/959 from 3 animals, Amyg-VTA: 0.21% or 1/479 from 2 animals). **G.** The experimental design for Retro-TRAP, top to bottom: Schematic of injection of retrogradely transported CAV-Cre in the Amyg and Cre-dependent AAV5 IV-GFPL10 virus in PL, followed by a 5-week expression period to allow for incorporation of GFPL10 fusion protein into ribosomes of Cre(+) neurons. The ribosomes tagged with GFP were then immunoprecipitated with GFP antibodies and the immunoprecipitated RNA was quantified. **H.** Significant differences were observed in enrichment levels of a panel of 10 marker genes between the 3 PL subpopulations (2-way ANOVA with neural subpopulation and gene as factors, p< .001, for main effect of subpopulation, and p < .001 for interaction of subpopulation X gene; n=3 groups for Amyg; n=3 groups for NAc, n=2 groups for VTA; each group is 6 mice).

To further confirm that PL-NAc, PL-Amyg, and PL-VTA neurons are distinct, we compared expression of a panel of 10 prefrontal cortical marker genes using the projection-specific molecular profiling technology Retro-TRAP [53,54] (Fig 1G and Supplementary Fig. 2). This revealed significant differences in enrichment levels in the panel of genes between the 3 subpopulations (Fig 1H; 2-way ANOVA with neural subpopulation and gene as factors, p< .001, for main effect of subpopulation, and p < 0.001 for interaction of subpopulation X gene).

### Activation of PL-NAc neurons, but not PL-VTA or PL-BLA neurons, decreases social investigation

Given that PL is composed of anatomically and molecularly distinct subpopulations that target the NAc, VTA and amygdala, we next asked if the previously reported effect of PL activation on social behavior [33] is broadly distributed across these descending projections, or if it can be localized to a specific projection. To address this question, we transiently activated either PL-NAc, PL-Amyg or PL-VTA neurons through ChR2 stimulation, while assessing social behavior using a 3-chamber test. Towards this end, an AAV5 virus expressing either ChR2-YFP or YFP-only (control mice) was injected into the PL (Fig. 2A), and optic fibers were implanted above the NAc, VTA or Amyg (Fig. 2B; fiber placement summary in Supplementary Fig 3). Before performing the 3-chamber test, the efficacy of the optogenetic terminal stimulation was verified with *in vivo* electrophysiology (Supplementary Fig. 4). During the behavioral test, mice explored a linear 3-chamber arena containing an encaged juvenile stranger mouse in one side-chamber (the “social target”) and an encaged novel object on the other side-chamber, with and without optogenetic stimulation (protocol: 3 minutes no-light, followed by 3 minutes of light, followed by 3 minutes no-light; Fig. 2C). Optogenetic stimulation of PL-NAc neurons resulted in a decrease in the amount of time mice spent near the social target, mimicking prior findings based on non-selective activation of PL neurons [33] (Fig. 2D; ANOVA with group, epoch, and light as factors; p=0.0196 for group X light). This decrease in social preference was mediated by a decrease in the duration of each social investigation bout (Supplementary Fig. 5A, p = 0.049, ANOVA group X light interaction), rather than an increase in the interbout interval (Supplementary Fig. 5B, p= 0.505, ANOVA group X light interaction). In contrast with the decrease in social preference, there was no decrease in novel object preference when PL-NAc neurons were stimulated, and in fact there was a trend towards increased novel object preference (Fig. 2G, p=0.579 for group X light).

**Fig. 2.**
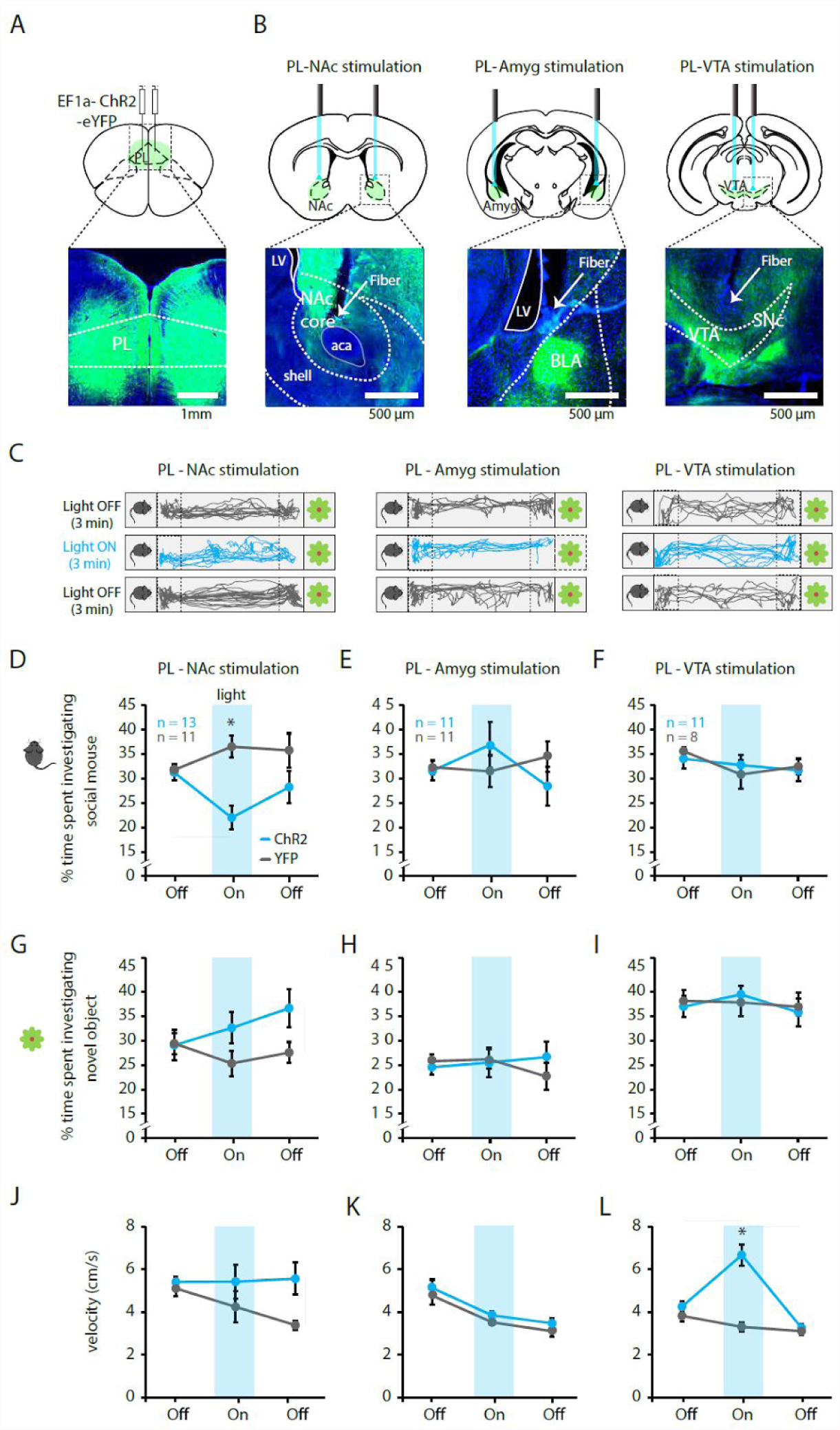
In the linear 3-chamber assay, activation of PL-NAc neurons, but not PL-VTA or PL-Amyg neurons, results in decreased social preference. **A.** A schematic (top panel) and coronal brain slice (bottom panel) of AAV5 CaMKii-ChR2-YFP injection in PL. **B.** Schematic (top panel) and histology (bottom panel) of fiber placement and terminal expression of ChR2-YFP in NAc, Amyg, and VTA. **C.** Position tracking from representative mice receiving activation of PL-NAc (left), PL-Amyg (center) or PL-VTA (right) neurons, while exploring the 3 chamber arena during pre-stimulation (top panel), stimulation (middle panel, 473 nm blue light, 20Hz stimulation, 5ms pulse duration, 7mW) and post-stimulation (bottom panel). Time spent near the social mouse or novel object (demarcated by dotted line) was measured. **D.** Stimulation of PL-NAc terminals decreased the amount of time mice spent in proximity of the social target (ANOVA with group, epoch, and light as factors; p=0.0196 for group X light). **E, F.** Stimulation of PL-Amyg (**E**) and PL-VTA (**F**) terminals had no effect on time spent in proximity of the social target (p=0.128 for group X light for PL-Amyg and p=0.406 for PL-VTA). **G-I.** Stimulating PL-NAc (**G**), PL-Amyg (**H**) and PL-VTA (**I**) terminals had no effect on time spent in proximity of the novel object (p=0.579 for Group x light for PL-NAc, p=0.625 for PL-Amyg and p=0.485 for PL-VTA). **J, K.** Stimulation of PL-NAc (**J**) and PL-Amyg (**K**) terminals had no effect on velocity (*p*=0.111 for group x light for PL-NAc, *p*=0.926 for PL-Amyg). **L.** Activation of PL-VTA neurons resulted in a large and significant increase in velocity (*p*<0.0001 for group x light). In panels **D-L**, error bars indicate SEM.

In contrast to stimulation of PL-NAc neurons, stimulation of PL-Amyg and PL-VTA neurons had no effect on social target preference (Fig. 2E,F; p=0.128 for group X light for PL-Amyg and p=0.406 for PL-VTA) or novel object preference (Fig. 2H,I; p=0.625 for group X light for PL-Amyg and p=0.485 for PL-VTA), pointing to a unique contribution of PL-NAc neural activity in supporting social preference.

To determine if the decrease in social preference mediated by PL-NAc stimulation could result from changes in locomotion, we compared the mouse’s velocity during the baseline and stimulation epochs in ChR2 and control mice. We did not observe a significant change in the velocity of the mice during stimulation of PL-NAc or PL-Amyg neurons, although there was a trend of increased velocity in the PL-NAc group in the post-stimulation epoch (Fig. 2J,K; p=0.111 for group X light for PL-NAc, p=0.926 for PL-Amyg). In contrast, activation of PL-VTA neurons resulted in a large and significant increase in velocity (Fig. 2L; p<0.0001 for group X light).

Taken together, PL-NAc stimulation caused a change in social preference but not in velocity, while PL-VTA stimulation caused a change in velocity but not social preference. This double dissociation between social investigation and velocity provides compelling evidence that a change in velocity alone cannot explain the decrease in social investigation resulting from PL-NAc stimulation. In addition, the similar density of PL-VTA, PL-NAc and PL-Amygdala neurons (Fig. 1D), and the similar proportions of downstream neurons affected by stimulation (Supplementary Fig. 4), suggests that the behavioral differences observed by stimulating these 3 projections is unlikely to result from differences in the efficacy of the stimulation.

Given that the 3-chamber test described above involves an encaged social target, we sought to determine if activation of PL-NAc terminals also reduced social investigation in a more natural assay of social investigation. In the home cage assay, a novel conspecific is introduced into the home cage of the test mouse and time spent in social investigation is scored (Fig. 3A). Using a counterbalanced experimental design, we found that PL-NAc activation (fiber placement summary in Supplementary Fig. 6) resulted in a large and significant reduction in social investigation in this assay as well, consistent with the reduction in social preference in the 3-chamber test (Fig. 3B,C, p=0.0001, paired t-test comparing the time spent investigating social target on control and stimulation days).

**Fig. 3.**
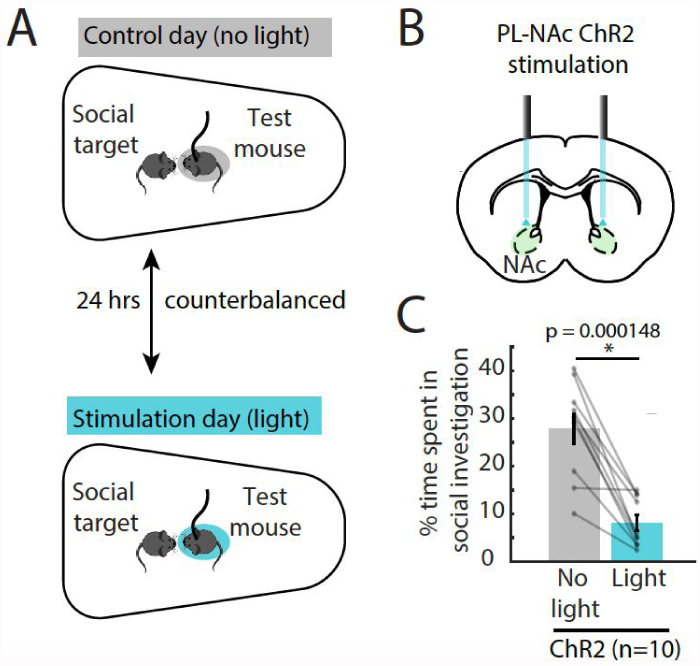
In the home cage assay, activation of PL-NAc neurons results in decreased social investigation. **A.** Schematic of the experimental design. Mice expressing ChR2 in the PL investigated novel social targets in their home cage across two days. A control day with no stimulation was counterbalanced with another day with stimulation (bottom panel, 473 nm blue light, 10 Hz stimulation, 5ms pulse duration, 1.5mW). Time spent investigating the social target was scored. **B.** Schematic of fiber placement targeting PL-NAc terminals expressing ChR2. **C.** Activation of PL-NAc terminals decreased the amount of time mice spent in social investigation (paired t-test; p=0.000148). Investigation was defined as epochs when the mouse was engaged in pursuit, sniffing, or grooming of the social target (see Methods).

### PL-NAc neurons are active during social investigation in a manner that depends on spatial location

Given that activation of PL-NAc neurons (and not the other subpopulations) decreases social preference, we sought to record the endogenous pattern of activity in these neurons during social behavior. To selectively record from PL-NAc neurons, we expressed the calcium indicator GCaMP6f selectively in PL-NAc neurons by injecting a retrogradely traveling CAV2 virus expressing Cre in the NAc and an AAV5 expressing Cre-dependent GCaMP6f in the PL (Fig. 4A). A head-mounted microscope [55] coupled to a GRIN lens was implanted above PL to obtain cellular-resolution images of PL-NAc neurons while mice explored the linear 3-chamber arena (Fig. 4A,B individual neurons were identified from videos using CNMFe [77]; Fig. 4C, example video in Supplementary Video 1;). A total of 830 neurons were imaged in 6 animals, and GRIN lens placement was confirmed post-hoc with histology (Supplementary Fig 7).

**Fig. 4.**
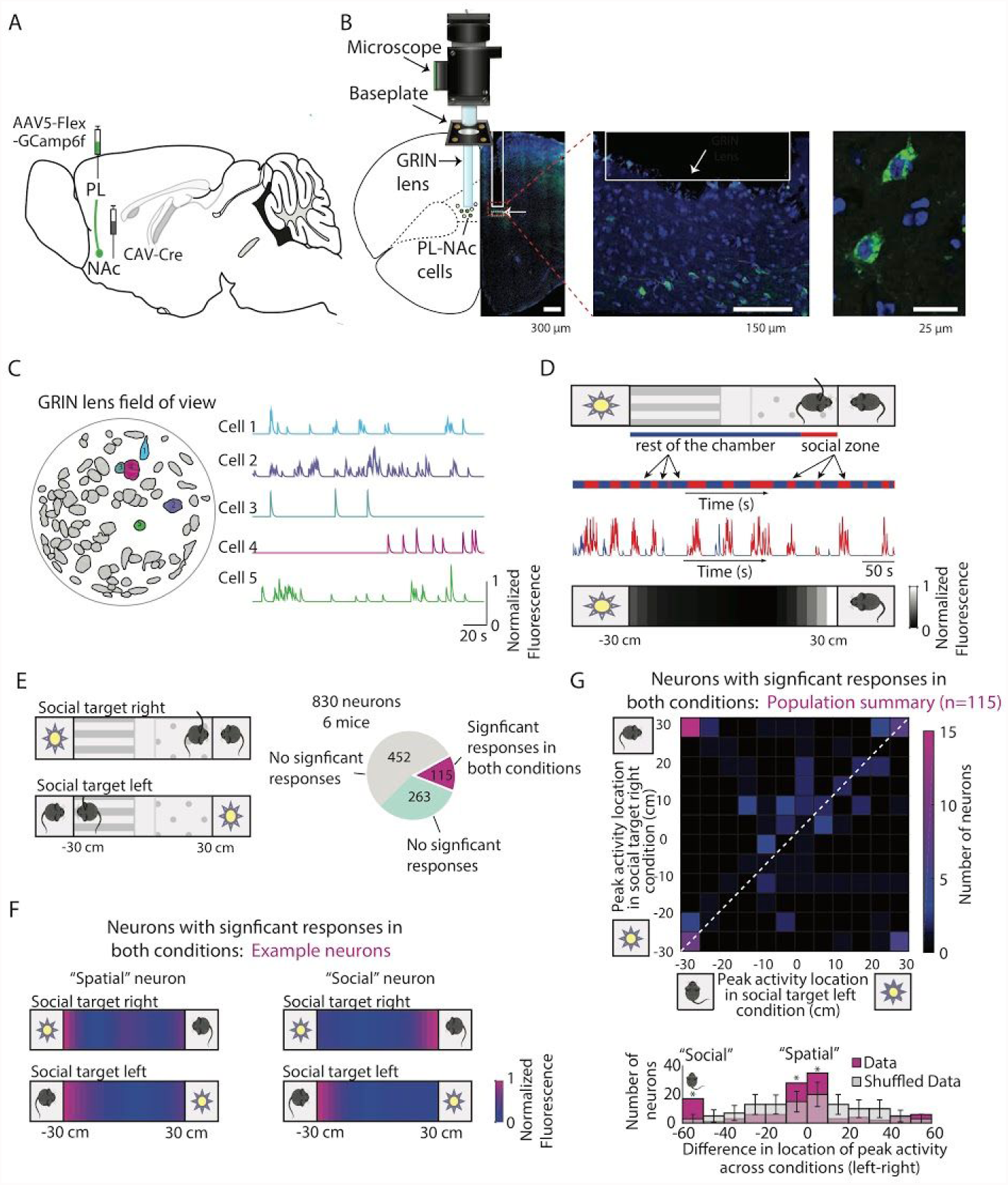
In the 3-chamber assay, some PL-NAc neurons are “spatial” and others are “social”. **A.** Targeting of gCaMP6f to PL-NAc neurons. Injection of retrograde CAV-Cre virus in the NAc and Cre-dependent AAV5 GCaMP6f in PL. **B.** Left: GCaMP6f imaging setup and histology of GRIN lens placement in PL. Middle: GCaMP6f expression in PL-NAc neurons in green and DAPI in blue. Scale bar: 150 μm. Right: A confocal image of PL-NAc neurons showing nuclear exclusion of GCaMP6f. Scale bar: 25 μm. **C.** The field of view of a GRIN lens in a representative animal with identified neurons demarcated (left panel). The right panel consists of fluorescence traces of example neurons (color coded in the left panel). **D.** Top panel: The imaged mouse explored a linear 3-chamber arena consisting of an encaged social target (stranger mouse) on one end and an encaged novel object on the other. Middle panel: Red denotes time that example mouse spent in “social zone” (10 cm from social target) while blue denotes other time. Normalized fluorescence of an example neuron showing elevated activity whenever the imaged mouse was in social zone. Bottom panel: Spatial receptive field of the example neuron. **E.** Mice were imaged while exploring the arena on two consecutive visits: first with the social target in one location (either “social target right” or “social target left”), and next with the social target in the other location. Of 830 neurons imaged from 6 mice, 14% (115 neurons) displayed significant responses in both conditions, while 31% (263 neurons) displayed significant responses in only 1 of the 2 conditions. The rest of this figure focuses on the 115 neurons with significant responses in both conditions. **F.** Left: The spatial field of an example “spatial neuron” that exhibited peak responses at a similar spatial location across both conditions (social target right condition on the top and social target left condition on the bottom). Right: The spatial field of an example “social neuron” that responded most strongly near the social target in both conditions. **G.** Population summary of neurons with significant responses across both conditions (n=115). Top: Number of neurons with peak response at each location across the two conditions. Bottom: Number of neurons as a function of the difference in location of peak activity across the 2 conditions reveals more “spatial neurons” and “social neurons” than expected by chance (p<0.001, 1 tailed t-test comparing data in each bin to simulated data generated from a spatially uniform peak distribution).

During exploration of the 3-chamber arena, we observed example neurons with elevated activity whenever the imaged mouse was in the vicinity of the social target (example neuron in Fig. 4D). However, since the social target is constrained to be in a specific spatial location in this assay, it is essential to distinguish potential social target-related responses from selectivity for the spatial location of the imaged mouse. To distinguish between these possibilities, all mice were imaged while exploring the arena on two consecutive visits (Fig. 4E): first with the social target in one location (either “social target right” or “social target left”), and next with the social target in the other location. 14% of neurons displayed significant spatial selectivity in both conditions, while 31% displayed significant spatial selectivity in only one of the two conditions (significance assessed based on cross-validated predictions of activity based on spatial receptive field estimates; see Methods).

Of the 115 neurons with significant selectivity in both conditions (“social target right” and “social target left”), some neurons exhibited peak responses at a similar spatial location across both conditions (example “spatial neuron” in Fig. 4F), while other neurons responded most strongly near the location of the social target in both conditions (example “social neuron” in Fig. 4F). In contrast, very few neurons responded consistently near the location of the novel object. In the population summary (top panel of Fig. 4G), neurons along the diagonal correspond to “spatial neurons” with the same spatial selectivity regardless of the location of the social target, while neurons in the upper left correspond to “social neurons” that respond in the vicinity of the social target, regardless of its spatial location. Across the population, there were more of these “spatial neurons” and “social neurons” than expected by chance (bottom panel in Fig. 4G, p<0.001, 1 tailed t-test comparing data in each bin to simulated data generated from a spatially uniform distribution of peak responses). Although the latter group of neurons provide a compelling neural correlate of social investigations, it is worth noting that they represent a relatively small fraction of the total imaged population (15 of 830, or 1.8%).

Interestingly, examining the population of neurons with significant responses in only one condition (“social target right” or “social target left”), rather than both conditions, revealed a substantially larger fraction of neurons that were active in the vicinity of the social target (90 of 830 neurons, or 10.8%, were most active in the spatial bin adjacent to social target, for one of the two conditions; Fig. 5). This is because many neurons exhibited elevated responses in the vicinity of the social target in only 1 of the 2 conditions, and showed no spatial selectivity at any location in the other condition (example neurons in Fig. 5B, population summary Fig. 5C and Supplementary Fig. 8). This phenomenon was observed for both the 1st and 2nd condition tested, implying that these responses cannot be explained based on greater social target novelty in the 1st condition (6.1% of all neurons responded most strongly near the social target in the 1st condition while 4.7% did so in the 2nd condition). In summary, many neurons responded in the proximity of the social target in a manner that depended on if it was in the left or right location. This implies that PL-NAc neurons encode the conjunction of social target proximity and spatial location.

**Fig. 5.**
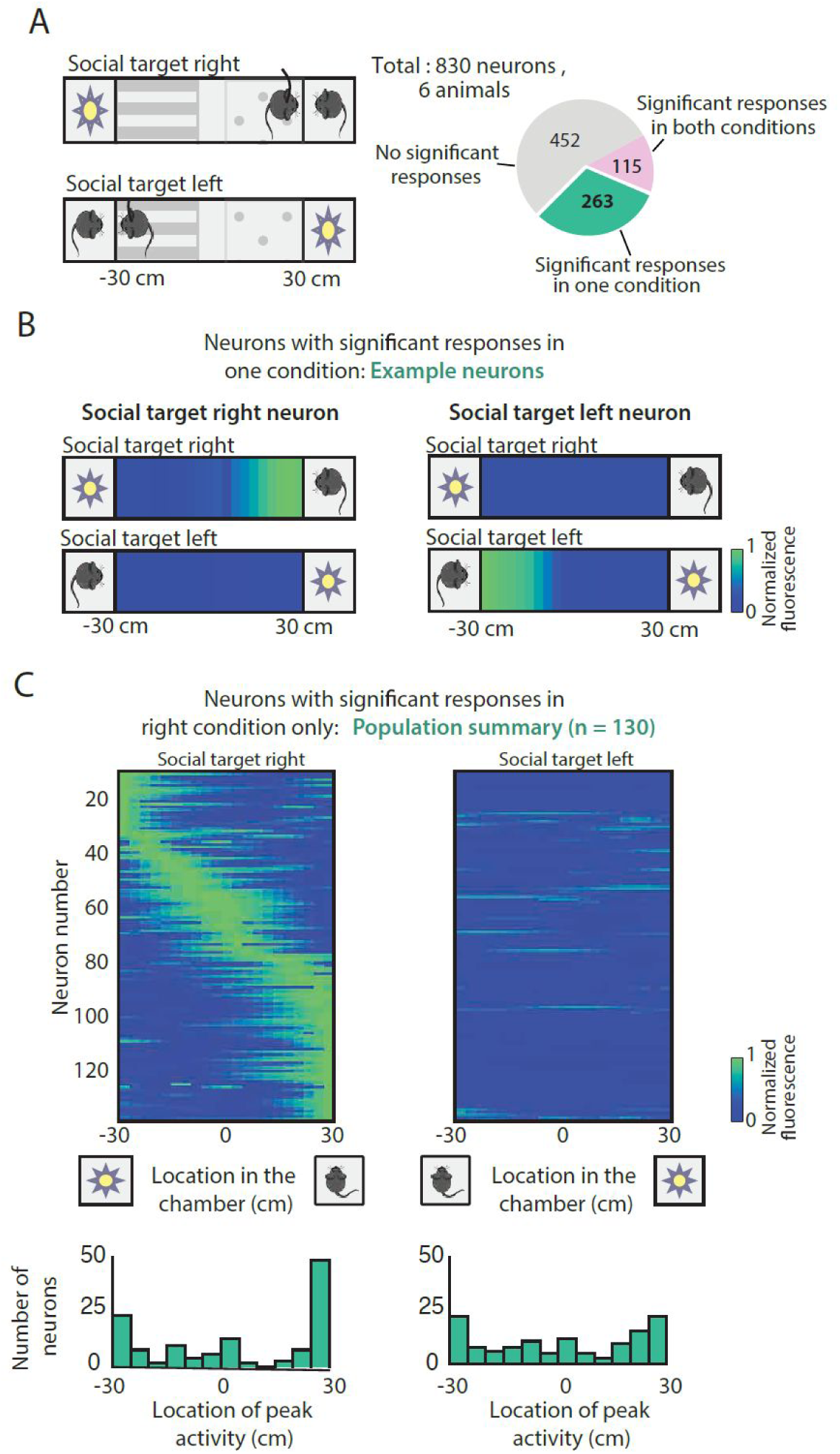
In the 3-chamber assay, many PL-NAc neurons that respond near the social target only do so in one spatial location. **A.** This figure focuses on the 263 (out of 830) PL-NAc neurons with significant responses in only one of the two conditions in the imaging experiment in the 3 chamber assay described in Fig. 4. **B.** Left panel: The spatial receptive field of an example neuron that exhibited spatial selectivity only in the social target right condition. Right panel: The spatial receptive field of a neuron that exhibited spatial selectivity only in social target left condition. **C.** Top panel: Spatial receptive fields of all neurons (n=130) with significant responses only in the social target right condition (left panel) and not the social target left condition (right panel). Bottom panel: Histogram of peak responses of the neurons along the length of the chamber. 38% of neurons responded most strongly in the vicinity of the social target, but only in one of the two spatial locations (in this case social target on the right).

Although the 3-chamber test provided a powerful approach to determine that social and spatial responses are combined in individual neurons, the encaged nature of the social target disrupts natural social interactions. Thus, to determine if PL-NAc neurons respond during social investigations between two freely moving mice, and if such correlates are modulated by spatial location as observed in the 3-chamber assay (Fig 4G, 5C), we imaged 243 PL-NAc neurons during the home cage assay across 3 mice (Fig. 6A, example video in Supplementary video 2). The locations of both mice were tracked and social investigation epochs were manually annotated (defined as periods when the imaging mouse was in pursuit, grooming or sniffing the social target). Some PL-NAc neurons, including the example neurons in Fig. 6B, responded during investigation of the social target (Fig. 6B).

**Fig. 6.**
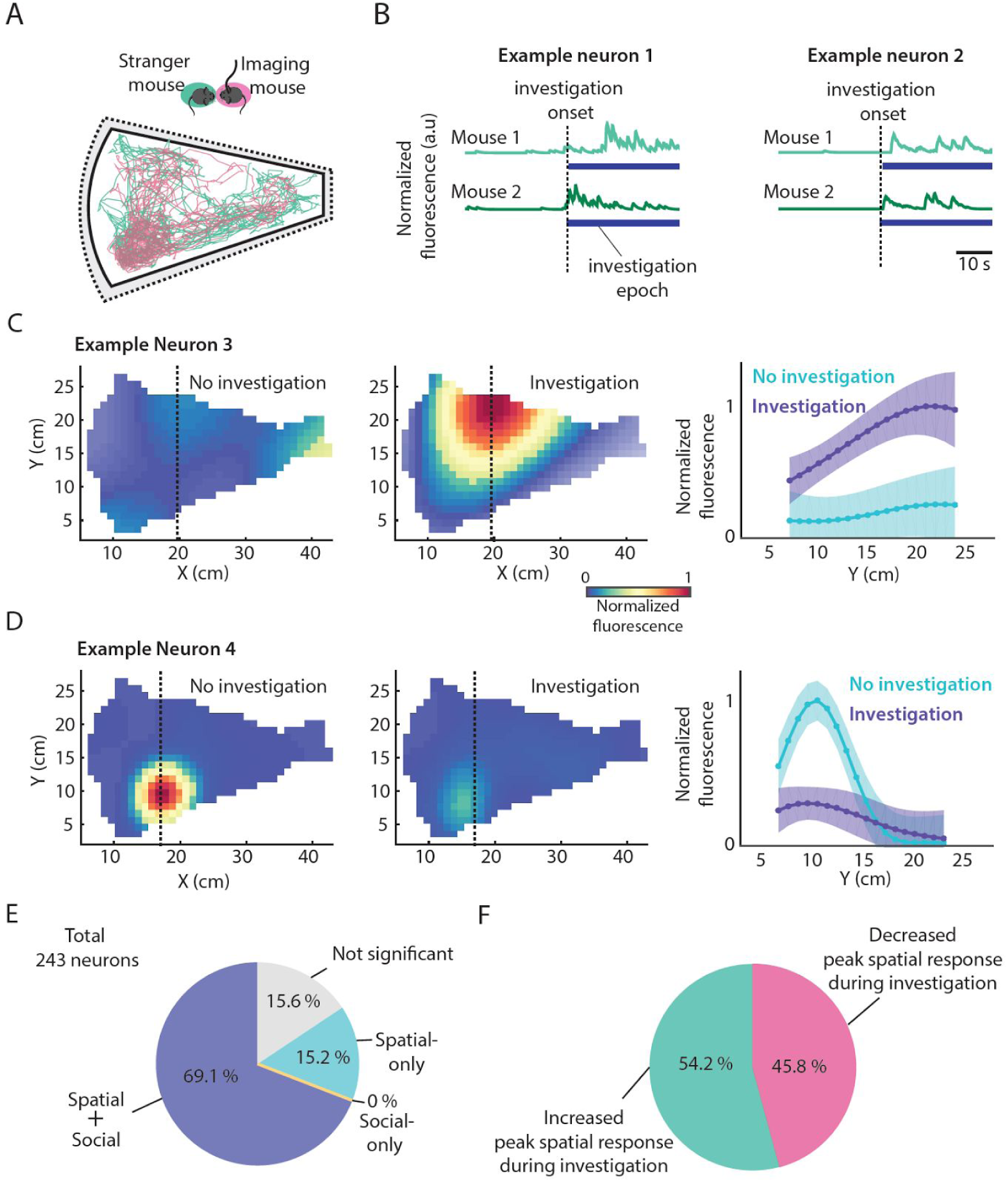
In the home-cage assay, PL-NAc neurons respond in a manner that depends on both social investigation and spatial location. **A.** Position tracking data of a representativ imaged mouse (pink) and a social target (green) during the home cage assay. Investigation was defined as epochs when the mouse was engaged in pursuit, sniffing, or grooming of the social target (see Methods) **B.** Example neurons with elevated activity during social investigation of multiple social targets (traces in green). **C.** An example neuron with spatial selectivity only during social investigation (right) but not in the absence of social investigation (left). Bottom panel: Traces of the estimated fluorescence from a slice (spatial bin with the maximum response) during investigation (purple) and no investigation (aqua). **D.** An example neuron with spatial selectivity only during epochs with no social investigation (left) but not during social investigation (right). Bottom panel: Traces showing the estimated fluorescence along a vertical slice (spatial bin with the maximum response) during social investigation (purple) and not investigation (aqua). Shaded regions denote error bars (2 standard deviations). **E.** Of the 243 imaged neurons, 69.1% had responses that were best explained by both social investigation and spatial position, while 15.2% were only spatial and 0% were only social. The remainder were not significantly modulated by either variable. These categories were determined based on comparing cross-validated fits of a social-only model, a spatial-only model, and a spatial+social model on cross-validated data (see Methods for details). **F.** Of the neurons with response modulation by both social investigation and spatial position (n=166), 54.2% increased their peak spatial response response during social investigation (as in the example neuron in **C**) while 45.8% decreased during investigation (example in **D**).

A striking result from the 3-chamber assay was that many PL-NAc neurons responded in the vicinity of the social target only in one of the two spatial locations that we tested (Fig 5). To determine if this combined spatial and social code generalized to the home cage assay, we took advantage of the fact that this assay involves bouts of social investigation interleaved with non-investigation epochs. This allowed us to calculate the spatial selectivity of each neuron during periods of investigation and also during periods without investigation. Comparison of spatial fields of individual neurons across these two conditions revealed modulation by social investigation: for example, some neurons exhibited spatial fields only during social investigation (example neuron in Fig. 6C) while others exhibited spatial fields only during epochs without social investigation (example neuron in Fig. 6D). To determine if this modulation of spatial receptive fields by social investigations was significant across the PL-NAc population, we compared the predictive power of two models on cross-validated data. One model was based on neurons having the same spatial receptive field regardless of whether or not the mice were interacting (spatial-only model), and the other model was based on calculating a different spatial receptive field during investigation versus during non-investigation epochs (social-spatial model). Across the population, the latter model produced significantly better predictions of neural activity in comparison to a model that only took into consideration spatial location (paired t-test comparing the correlation coefficients of the predicted and real data across the two models, *p*= 6.23e-22, *n*=243 neurons). In addition, 69.1% of neurons had activity that was best predicted by the social-spatial model while only 15.2% had activity best predicted by spatial-only model (Fig. 6E). Of neurons best described by the social-spatial model, similar fractions increased versus decreased their peak spatial response during investigation (Fig. 6F). In summary, these findings demonstrate that PL-NAc neurons encode the conjunction of social and spatial information in the home cage assay.

### Spatially-specific inhibition of PL-NAc neurons disrupts social-spatial learning

What could be the function of a combined social-spatial code in PL-NAc neurons? Given that plasticity of PL synapses in NAc is thought to be a site of reward-related learning [56–61], we hypothesized that these these neurons could support the formation of social-spatial associations by providing information to the NAc about the location of social encounters. In order to test this hypothesis, we designed a specialized social conditioned place preference paradigm which involved a test mouse exploring two chambers, each with an encaged social target. This paradigm enabled us to differentially modulate the formation of a social preference for one location but not the other (Fig. 7C). To selectively inhibit PL-NAc neurons, we used an intersectional virus strategy to express the inhibitory opsin eNpHR3.0-eYFP (or control virus; Fig 7A-B). Before conditioning, mice first experienced a “baseline day” in which there was no social target and no optogenetic stimulation, in order to ensure the mice exhibited no initial spatial bias. They then received two conditioning days in which the neurons were inhibited in the vicinity of one social target but not the other, followed by a final test day to assess persistent changes in spatial preference (Fig. 7C). We found that mice that received inhibition of PL-NAc neurons did not form an association with the social zone associated with inhibition (relative to control mice) (Fig. 7D, ANOVA with group and days as factors; p=0.0271 for group X days, followed by post-hoc t-test Baseline: p=0.378, Cond day 1: p=0.008, Cond day 2: p=0.025, Test day: p= 0.024), while the preference for the social zone without inhibition was indistinguishable from controls (Fig. 7E, ANOVA with group and days as factors; p=0.681). This lack of preference for the social zone associated with inhibition persisted to the next day, suggesting that activity in PL-NAc neurons is necessary for forming a social-spatial preference.

**Fig. 7.**
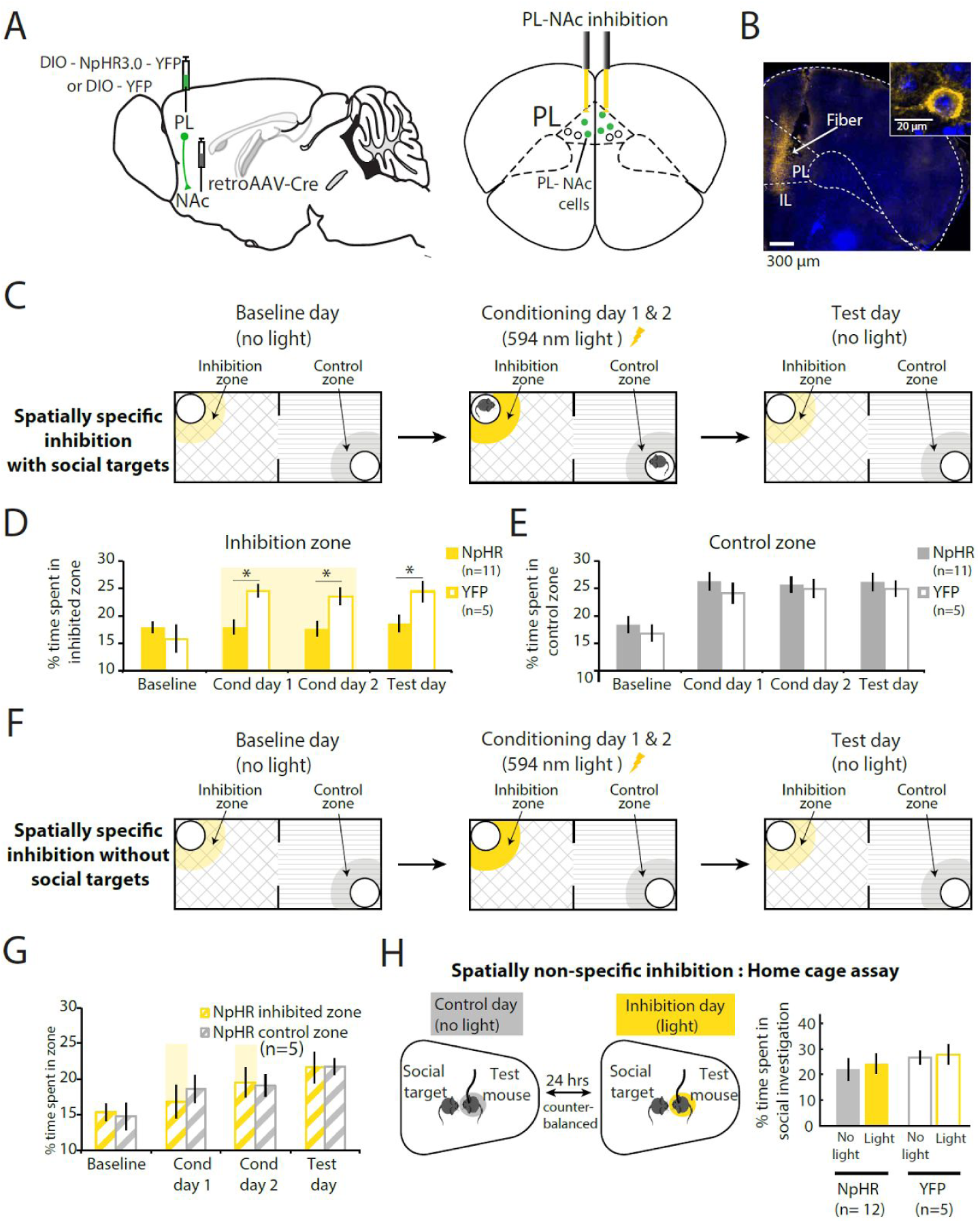
Spatially-specific inhibition of PL-NAc neurons disrupts social-spatial learning. **A.** Left: To target NpHR expression to PL-NAc neurons, the retrograde virus retroAAV-Cre was injected in the NAc and Cre-dependent AAV5-DIO-NpHR or AAV5-DIO-YFP was injected in PL. Right: Schematic of fiber placement in PL. **B.** Histology of fiber placement and terminal expression of NpHR-YFP in PL-NAc neurons (YFP expression in yellow and DAPI in blue). **C**. Social conditioned place preference paradigm with spatially specific inhibition. Baseline day: Mice explored the two-chamber arena with empty cages at two ends. Conditioning days (2 days): Mice explored the same two-chamber arena, each with an encaged social target. PL-NAc neurons were inhibited in the proximity of one social target (594 nm, 2.5-3mW) but not the other. Test day: Mice explored the arena in the absence of both social targets and inhibition. Each day the time spent near the cages (social zone) was measured. **D**. In contrast to control mice (YFP, n=5 animals), mice expressing NpHR (n=11 animals) in PL-NAc neurons failed to form a preference for the social zone paired with yellow light (conditioning days) and this effect persists the day after (test day) in the absence of any inhibition (ANOVA with group and days as factors; p=0.0271 for group X days, followed by post-hoc t-test Baseline: p=0.378, Cond day 1: p=0.008, Cond day 2: p=0.025, Test day: p= 0.024). **E.** Both control and YFP mice acquired a preference for the social zone (control zone) that was not paired with light (ANOVA with group and days as factors; p=0.681). **F.** The conditioning paradigm was performed as above (7C) but in the absence of any social targets. **G.** In the absence of social targets, inhibition of PL-NAc neurons had no effect on spatial preference (ANOVA with zones and days as factors; p=0.775). **H.** Left: Schematic of the home cage assay. Right: Spatially nonspecific inhibition of PL-NAc neurons had no effect on the time mice spent in social investigation (ANOVA with group and days as factors; p=0.825).

Although these results are consistent with with the idea that PL-NAc neurons support the formation of social-spatial associations, there are other possibilities that must be examined before that conclusion can be definitive. First, it is possible that inhibition of PL-NAc neurons is aversive, and therefore inhibition of those neurons would result in altered spatial preference in the conditioning assay described above (Fig. 7C), even in the absence of a social target. To examine this possibility, in a new group of mice we repeated the same conditioning paradigm as described above, inhibiting the PL-NAc neurons in the same zone of the same chamber, but critically, in this case the social targets were omitted (Fig. 7F). In this case, inhibition had no effect on spatial preference (Fig. 7G, ANOVA with zones and days as factors; p=0.775), supporting the conclusion that inhibition of PL-NAc neurons does indeed disrupt social-spatial learning.

In addition, it is also important to control for the possibility that inhibition of PL-NAc neurons decreases an animal’s preference for social investigation behavior itself. In this view, PL-NAc neurons could contribute to the subjective value of social investigation, or to physically performing the behavior, rather than to the formation of social-spatial associations. If this were the case, we would expect that spatially nonspecific inhibition of PL-NAc neurons during a home cage social assay, which involves two freely moving mice, would result in decreased social investigation. In fact, we observed no change in social investigation in the home cage assay as a result of spatially nonspecific inhibition of PL-NAc neurons (Fig. 7H, ANOVA with group and days as factors; p=0.825). Thus, together these results suggest that PL-NAc neurons contribute to social-spatial learning.

## Discussion

Here, we show that PL is composed of distinct populations of neurons that target 3 regions that have been implicated in social behavior: the NAc, VTA, and amygdala. Activation of PL-NAc neurons, but not the other descending projections from PL, decreases social investigation across two behavioral assays. In both assays, social responses in PL-NAc neurons depend on spatial location. To test the hypothesis that combined social-spatial coding in these neurons could enable the formation of social-spatial associations, we inhibited this population during social investigation only in a specific spatial location. Consistent with our hypothesis, spatially-specific inhibition prevents the formation of a preference for the location of the social encounter, while spatially non-specific inhibition has no effect on social investigation. These results suggest that the unexpected combination of spatial and social information within the PL-NAc population supports socially motivated behavior by enabling the formation of social-spatial associations. This mechanism may reflect a very general and fundamental contribution of this projection to reward-context learning, even beyond social behavior.

### Relevance of a combined social-spatial code in PL-NAc neurons

The combined social and spatial code that we observed in PL-NAc could provide a powerful template to support learning to perform appropriate socially motivated behavior in a location-specific manner. In theory, such learning could be mediated by dopamine-dependent synaptic plasticity in the NAc. Prefrontal cortical inputs to the NAc are thought to be strengthened when activated in concert with dopamine release [62]. In this view, PL-NAc synapses that are active during social investigation in a specific spatial location would be strengthened, given the release of dopamine that is known to accompany social investigation (Fig. 8A) [26]. Such plasticity could in turn reinforce conditioned approach behavior to the location of the encounter (i.e. a social-spatial association).

**Fig. 8.**
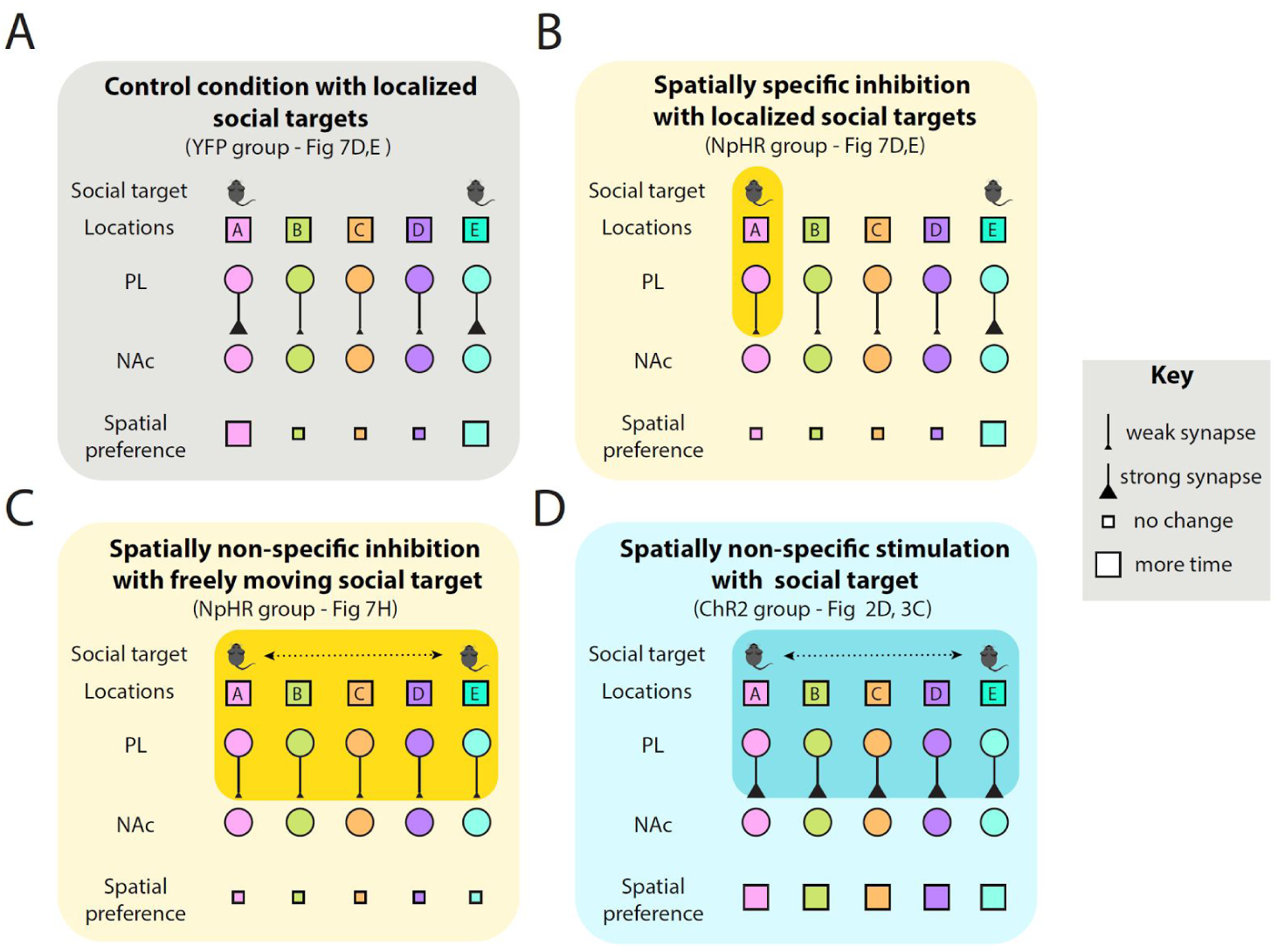
Schematic of plasticity mechanisms that could relate optogenetic manipulation of PL-NAc neurons to social behavior. In all panels, PL-NAc neurons are active during social investigation in a specific location (location A, B, C, D, or E). The strength of the synaptic connection between a PL neuron and its target in NAc increases during social investigation for the PL neuron that encodes that location. The strength of the PL-NAc synapse determines the mouse’s spatial preference for the corresponding spatial location. **A.** In this case (YFP control group in Fig 7D,E), there is a target mouse in location A and E. Therefore, synaptic connections of PL-NAc neurons that respond during social investigation in those locations are strengthened, leading to an increase in preference for both location A and E. **B.** Same as panel **A**, but the PL-NAc neurons that respond in location A are silenced (NpHR group from Fig 7D,E). In this case, synaptic connection of location A neurons are not strengthened, but location E neurons are strengthened. **C.** During freely moving behavior (Fig. 7H), there is no change in synaptic strength during social investigation, since it does not occur in a specific spatial location. Therefore, inhibition does not prevent a change in synaptic strength. **D.** Spatially nonspecific excitation may instead strengthen PL-NAc synapses, regardless of location (Fig 2D, 3C). This may effectively increase preference for non-social locations, not only social locations, effectively decreasing preference for the social location. This non-specific synaptic strengthening may also explain the trend towards decreased social investigation after the optical stimulation.

Consistent with this hypothesis, spatially specific inhibition of PL-NAc neurons in the location of a social encounter prevented the formation of a social-spatial association for that location (Fig. 8B). This manipulation should preferentially silence PL-NAc neurons that are normally active during social investigation in the inhibited location, and may therefore prevent the synaptic weights of those neurons from increasing in the NAc, while allowing synaptic weights of other neurons to increase (e.g. the weights of synapses that are active during investigation of the “control” social target,;Fig 7E). In contrast, spatially uniform inhibition would silence all PL-NAc neurons equally (Fig. 7H), and therefore no synaptic weights would be expected to change during the period of stimulation (Fig. 8C). The lack of consequence of spatially uniform inhibition of PL-NAc neurons suggests a function for PL-NAc neurons in social-spatial learning, rather than in encoding the value of social interactions, or in the performance of social investigation.

Why might spatially uniform activation have reduced social investigation (Fig. 2D, 3C), given that spatially uniform inhibition did not have this effect (Fig. 7H)? Continuous activation of PL-NAc neurons might uniformly strengthen the synaptic weights of those neurons in NAc, regardless of the spatial location that the neurons encode (Fig. 8D). Uniform strengthening of PL-NAc synapses may reduce the relative preference for the location of the social target by effectively increasing preference for other locations. Such nonspecific synaptic strengthening as a result of continuous PL-NAc activation could also explain behavior in the post-stimulation epoch, when there was a trend towards a sustained reduction in social investigation and increase in novel object investigation (Fig 2D,G; Supp Fig 5).

### Conjunctive codes in prefrontal cortex

The combined coding of social and spatial information in PL-NAc neurons is reminiscent of highly conjunctive codes that have been reported in non-human primate prefrontal cortex in decision making tasks [63,64]. It has previously been suggested that the computational advantage of such “mixed selectivity” is to enable flexible readout of parameters of interest by a linear decoder.

Here, we propose an additional and complementary function of conjunctive codes. As described in detail above, we propose that a conjunctive code for spatial position and social investigation would provide information about an animal’s current state, which is an essential ingredient for reinforcement learning. Thus, a reinforcement learning algorithm (e.g. Q-learning) could update the value of a spatial location, as represented in NAc, by combining social-spatial information in PL-NAc neurons with a reward prediction error carried by dopamine neurons [65-68].

Through analogous mechanisms, a conjunctive code in PL-NAc neurons may potentially also support learning about other reward-location associations, beyond social-spatial learning. For example, PL-NAc neurons may also respond during consumption of food reward, with different neurons selective for different spatial locations, a coding scheme which could potentially support food-context learning. In fact, PL-NAc neurons have been previously implicated in reward-seeking for food and drug reward [56,57,59,69–71], although to our knowledge a contribution to the spatial aspect of such behaviors has not been investigated.

The conclusion of combined social-spatial coding depended on rigorous analysis of the relationship between neural activity and behavior, which is difficult in the context of social behavior, in part because of the relatively limited amount of time mice spend engaged in social interactions. To overcome this challenge, we employed a Bayesian framework based on Gaussian processes to accurately estimate the relationship between neural activity and behavior (see Methods).

### Connections to previous research on social behavior and remaining open questions

An important question had been why activation of PL caused mice to spend less time investigating a conspecific, when (spatially non-specific) inhibition of endogenous activity resulted in no observable change in behavior [33]. Here, we resolved this question by demonstrating that under the right conditions, inhibition of endogenous activity in PL-NAc neurons does disrupt social behavior. Specifically, spatially-specific inhibition disrupts the formation of a social-spatial preference, supporting a contribution to social-spatial learning. Given that NAc has been implicated in social learning [27–30,46], and given that social investigation results in dopamine release from the VTA into the NAc [26], our results provide a coherent link between previously disparate literatures regarding the involvement of PL, NAc and VTA in social behavior.

A remaining open question is which anatomical inputs provide spatial information to PL-NAc neurons. An obvious candidate is the ventral hippocampus, a region well known for its spatial coding, and which projects directly onto PL-NAc neurons [57,72]. A related question is which inputs provide social information to PL-NAc neurons. Visual, olfactory, somatosensory, and auditory inputs could potentially contribute to the observed neural responses. One possibility is that a reduced social stimulus comprised of only one of those modalities is sufficient to generate social responses in PL-NAc neurons; alternatively, the responses may fundamentally require a multimodal social target. Yet another interesting question is if the responses are better described as sensory or motor. The former would correspond to responses that are better predicted by the behavior of the target mouse, while the latter would correspond to responses that are better predicted by the imaged mouse’s behavior. Dissociating between these possibilities may ultimately be possible through detailed measurements and modeling of the behavior of both the imaged as well as the target mouse.

### Conclusions

In conclusion, we found that a combination of social and spatial information within PL-NAc neurons supports socially motivated behavior by enabling the formation of social-spatial associations. This represents a novel mechanism through which PL contributes to social behavior, which may generalize to other forms of reward-location learning.

**Supplementary video 1.** Representative movie demonstrating PL-NAc neuron activity, related to Figure 4, 5 and 6.

Raw grayscale movie (motion corrected and cropped) showing isolated PL-NAc neuron activity (in white). Video sped up 4 times.

**Supplementary video 2.** An example movie showing PL-NAc neuron activity during social investigation in the home cage assay, related to Figure 6.

The behavioral video on the left and the related related PL-NAc imaging data on the right (motion corrected-cropped grayscale movie). Videos sped up 4 times.

## Materials and Methods

### Animals

All experimental procedures were conducted in accordance with the National Institutes of Health guidelines and were reviewed by the Princeton University Institutional Animal Care and Use Committee (IACUC). Male C57BL/6J mice were used for these experiments. Mice were maintained on a 12-hour light on – 12-hour light off schedule. All surgical manipulations and behavioral tests were conducted during their light off period.

### Stereotactic surgeries

Mice were anaesthetized with 1-2% isoflurane and placed in a stereotactic setup. A microsyringe was used to deliver virus or neural tracer to target brain regions.

For retrograde tracing experiments, 0.3 – 0.5 μL of cholera toxin β subunit conjugated to fluorophores (CTB-488, CTB-555, CTB-647m, Life Technologies Corporation) were injected unilaterally into the NAc core (1.3 mm anterior, 1.0 mm lateral and 4.7 mm in depth, n=6 animals), VTA (3.3 mm posterior, 0.7 mm lateral and 4.7 mm depth, n=7 animals) and amygdala (1.6 mm posterior, 2.8 mm lateral and 4.8 mm depth, n=5 animals) of ~6-10 week old male mice.

For molecular profiling experiments, to specifically target PL projection subpopulations, ~2 month old male mice were bilaterally injected in the PL (1.8 mm anterior, 0.5 mm lateral and 2.5 mm in depth) with 1μL of AAV5-IV-GFPL10 and 0.75 μL of a retrogradely transporting CAV-Cre virus (IGMM Vector core, France, ~2.5 × 1012 parts/ml) into the Amyg, NAc and VTA.

For optogenetic activation experiments, ~1.5 month old male mice were bilaterally injected in the PL (1.8 mm anterior, 0.5 mm lateral and 2.5 mm in depth) with 1 μL of AAV5-CamKIIa-hChR2-EYFP (n = 35 ChR2 mice, UNC Vector Core, 4 × 10_12_ parts/ml and 8 × 10_12_ parts/ml) or AAV5-CamKIIa-EYFP (n = 30 control mice, UNC Vector Core, final titer of 7.5 × 10_12_ parts/ml) virus. Ferrules attached to optical fibers (300 μm core diameter, 0.37 NA) were implanted bilaterally to target either the NAc core (1.3 mm anterior, 1.0 mm lateral and 4.3 mm in depth, ChR2 n=13, YFP n=11 animals, Home cage assay, ChR2 n=10), Amyg (1.6 mm posterior, 3.4 mm lateral and 4.3 mm in depth, ChR2 n=11, YFP n=11 animals) or the VTA (3.3 mm posterior, 0.7 mm lateral and 4.3 mm depth, ChR2 n=11, YFP n=8 animals).

For imaging experiments, to specifically target PL neurons projecting to the NAc, ~ 2 month old male mice (n=9 animals) were injected with 0.75 μL of a retrogradely transporting CAV-Cre virus (IGMM Vector core, France, ~2.5 × 1012 parts/ml) into the NAc (1.3 mm anterior, 1.0 mm lateral and 4.7 mm in depth) and 0.75 μL of AAV5-CAG-Flex-GCaMP6f-WPRE-SV40 (UPenn Vector Core, injected titer of 3.53 × 1012 parts/ml) into the PL (1.8 mm anterior, 0.5 mm lateral and 2.5 mm in depth). A week post injection, mice were implanted with a 0.5mm diameter, ~6.1mm length GRIN lens (GLP-0561, Inscopix) or a 1 mm diameter, ~4.3 mm length Prism probes (PPL-1043, Inscopix) over the PL (1.8 mm anterior, 0.3 mm lateral and 1.9-2.0 mm in depth). A plastic cap was glued to protect the lens. Mice were housed individually post lens implant. 3-4 weeks after the injection, a baseplate (BPL-2, Inscopix) attached to the miniature microscope (nVISTA HD v2, Inscopix) was positioned over the GRIN lens such that the neurons and/or other structures (such as blood vessels) in the brain were in focus. The baseplate was then implanted using dental cement, and a baseplate cover (BPC-2, Inscopix) protected the GRIN lens.

For optogenetic inhibition experiments, to specifically target PL neurons projecting to the NAc, ~ 1.5 month old male mice (NpHR: n=12 animals, YFP: n=5 animals) were injected with 0.75 μL of a retrogradely transporting retroAAV-Cre virus [73] (PNI Viral core, 1.16 x 10^14^ parts/ml) into the NAc (1.3 mm anterior, 1.0 mm lateral and 4.7 mm in depth) and 0.5 μL of AAV5-EF1a-DIO-NpHR3.0-EYFP (UPenn Vector Core, injected titer of 1.29x10^13^ parts/ml) or AAV5-EF1a-DIO-YFP (UPenn Vector Core, 8.41x10^12^ parts/ml) into the PL (1.8 mm anterior, 0.5 mm lateral and 2.5 mm in depth). Ferrules attached to optical fibers (300 μm core diameter, 0.37 NA) were implanted bilaterally to target the PL.

### Immunohistochemistry

Mice were anesthetized with 0.08 ml Euthasol (i.p. injection) and transcardially perfused with 1x phosphate-buffered saline (PBS), followed by fixation with 4% paraformaldehyde in PBS (PFA). Brains were dissected out and post-fixed in 4% PFA overnight before being transferred into 30% sucrose in PBS solution. 40 μm thick coronal sections containing the brain region of interest were cut on a freezing microtome.

For the retrograde tracing experiments, we first verified that the CTB injections were localized to the target brain regions (NAc, VTA and the Amyg). Coronal sections containing the PL were then imaged (Nikon Ti2000E) in order to confirm retrograde labeling of NAc-projecting, Amygdala-projecting, and/or VTA-projecting neurons in PL. Using The Mouse Brain in Stereotaxic Coordinates, Second Edition, by George Paxinos and Keith B. J. Franklin (2001), as a reference, for each animal three CTB-labeled brain sections were chosen along the anterior-posterior axis (~1.5mm, ~2.2 mm, and ~2.6 mm anterior to bregma). Cellular resolution images of coronal sections of interest were then acquired through the 63x immersion objective using the Leica TCS SP8 confocal microscope. The boundary of prelimbic cortex was determined using measurements from a mouse brain atlas [74], and all CTB-labeled cells within PL were assessed. The extent of co-labeling and the distance from the midline were measured for each cell using tools in the Leica imaging software suite. For the data presented in Fig 1E, the M/L cell distributions were averaged at 3 anterior-posterior locations (1.5 mm, 2.2 mm and 2.6 anterior to bregma, for separate M /L cell distributions at 3 A/P locations see Supplementary Fig. 1D-L). 12/30 mice were excluded from the CTB experiments because of failed injection or for not showing any CTB expression.

For the optogenetic activation experiments, a GFP antibody stain was used to enhance visualization of virus expression in terminals. The primary antibody was a mouse monoclonal anti-GFP (1:1,000, Life Technologies, No. G10362) and the secondary was a donkey anti-rabbit coupled to Alexa 488 (1:1,000 dilution, Jackson ImmunoResearch, No. 711-545-152). Mounted sections were imaged to confirm expression of the virus in PL cell bodies and terminals in the NAc, VTA and the Amyg and optical fiber targeting in the downstream structures. 9/74 mice were excluded from the optogenetic studies, 2 mice for seizing during stimulation, 6 mice for failed fiber targeting and 1 mouse for technical problems encountered during behavioral testing.

For the GCaMP6f imaging experiments, coronal sections of PL were used to confirm virus expression, nuclear exclusion of GCaMP6f, and to verify targeting of the GRIN lens to PL. 5/14 mice were excluded from the imaging experiments, 1 mouse for GRIN lens implant being too posterior and 4 mice for showing no GCaMP6f expression.

For the optogenetic inhibition experiments, coronal sections of PL were used to confirm virus expression, nuclear exclusion of NpHR, and to verify targeting of the fibers to PL.

### Molecular profiling experiments

Approximately 5 weeks post-injection, mice were sacrificed and their brains were rapidly dissected on ice to isolate medial prefrontal cortex. The dissected brain tissue from 6 mice were combined to form a single sample (number of samples-Amyg: 3, NAc: 3, VTA: 2). As described previously [53,54], GFPL10-positive ribosomes were then immunoprecipitated using a 2 hr GFP immunoprecipitation (GFP IP). qPCR analysis was performed using TaqMan assays for the targeted genes to determine differential enrichment. Fold-enrichment was calculated as IP RNA/Total RNA. Expression was compared across the subpopulations with a 2-way ANOVA with neural subpopulation and genes as factors.

### *In vivo* electrophysiology experiments

~ 15 weeks after injection of ChR2 in PL, mice were anaesthetized with urethane (1.2mg/g). A ~125 μ m tungsten electrode (A-M systems) was glued to a fiber optic (300 μm core diameter) to make an optrode. The optrode was then lowered to the recording sites using stereotactic coordinates (PL:1.8 mm anterior, 0.5 mm lateral and 2.5 mm in depth, NAc: 1.3 mm anterior, 1.0 mm lateral and 4.3 mm in depth, Amyg:1.6 mm posterior, 3.4 mm lateral and 4.3 mm in depth or the VTA:3.3 mm posterior, 0.7 mm lateral and 4.3 mm depth). Spike 2 (CED) was used for data acquisition. To verify ChR2 expression in PL and its terminals in NAc, VTA and BLA, light pulses of 2 seconds (447nm, 5 mW at the tip of the optical fiber) were delivered at regular intervals (every 15 seconds) for many trials (30-50). Spike 2 in combination with custom MATLAB software was used for analysing the data. A paired t-test comparing firing rates of individual neurons before and during light stimulation was used to determine if a neuron was significantly modulated by light stimulation.

### Behavior Assays

Linear 3-chamber assay: In this assay, mice explored a linear three-chamber arena with distinct textured floors containing an encaged juvenile stranger mouse in one side-chamber (the “social target”) and an encaged novel object on the other side-chamber. Ethovision (Noldus) was used to track the location of the mouse for all linear 3-chamber assays.

Homecage assay: On the day of the assay, the imaging mouse was allowed to explore his homecage for 30 minutes, following which an unfamiliar juvenile male mouse (~6 weeks, “social target”) was introduced into his homecage for a 5 min period. All behavior was video monitored. Social investigation was defined as periods when the imaging mouse was in pursuit (defined as the period from when the animal orients and runs towards the social target to the point when he makes contact), sniffing (defined as periods when the mouse’s snout was in contact with the social target) or grooming (as periods when the forepaws of the mouse and the mouth and snout region were placed gently on the social target for licking) the social target. Investigations of the social target by the imaging mouse were manually annotated (using JABBA, Janelia). For a subset of the experiments, the location of both the imaging mouse and the social target was tracked post hoc (in a semi-automated fashion using Adobe After Effects).

For optogenetic experiments, mice were allowed to investigate a novel social target introduced into his homecage for a 5 minute period. The assay was repeated again after a 24 hour interval with a another novel social target. The amount of time spent investigating the social target was manually annotated.

Social conditioned place preference paradigm: In this assay, on baseline day mice explored a two-chamber arena containing two empty cages at each end. Distinct walls and floors distinguished the two chambers. This was followed by 2 days of conditioning, mice were explore the arena with each chamber containing an encaged social target. On the test day mice were allowed to explore the arena with. A 10 cm arc surrounding the cages was designated as the social zone and time spent in each social zone was measured each day.

In a subset of the animals (NpHR n=5), this assay was carried out as above but with no social targets in the cages on all days (baseline, conditioning and test day). These mice (4/5 mice, one mouse was excluded for leaving the arena multiple times during behavioral testing) were rerun on the social version of this assay a week later after establishing the absence of any spatial bias for either zone on the baseline day.

### Optogenetic activation of ChR2 expressing PL terminals

For optogenetic experiments, ~14 weeks after injection of ChR2/YFP virus in PL, mice were run on the linear 3-chamber assay. The experimenter was blind to the treatment group during the course of the assay. The mice were allowed to explore the linear chamber for 3 min with no light stimulation, followed by 3 min of blue light stimulation (447nm, 7-10 mW at the tip of the optical fiber; 5 ms long pulses at 20Hz) and terminating with an additional 3 min of no light. The location of the mice was tracked using Ethovision (Nodulus).

A multifactor ANOVA, consisting of three factors: groups (ChR2, YFP), light (OFF, ON, OFF) and time epochs (first 3 minutes, middle 3 minutes, last 3 minutes) followed by a Tukey corrected post-hoc tests to determine significance.

For the other the stimulation experiment, mice expressing ChR2 in PL-NAc terminals (n=10) were run on the homecage assay. On one of two days designated as the stimulation day, the mice were allowed to investigate a social target introduced into their homecage while they were stimulated with blue light (447 nm, 1.5 mW, 5ms long pulses at 10Hz). On the control day, mice explored investigated a novel social target while tethered but receiving no light stimulation. Half the mice received stimulation on day 1 while the rest on day 2.

### Optogenetic inhibition of NpHR expressing PL-NAc neurons

~ 4 weeks post injection mice expressing NpHR/YFP in PL-NAc neurons were run on the Social conditioned place preference paradigm. For each animal one social zone was designated as the control zone and the other zone the inhibited zone (in random). On the conditioning days, the inhibited zone was paired with yellow light stimulation (594nm, 2.5-3 mW, constant light). Light remained on when the animal moved into this zone and remained on until the animal moved out of the zone. Mice received no stimulation in the control social zone. On the test day, mice were allowed to explore the arena in the absence of light stimulation and in the absence of social targets to determine their preference for the social zones. A multifactor ANOVA, consisting of two factors: groups (ChR2, YFP) and days (Baseline, Conditioning day 1, Conditioning day2, Test day) followed by a post hoc t-tests comparing time spent in the each zone by the control and NpHR animals was used to determine significance.

For a subset of the NpHR mice (n=5 animals), the assay was run as above but in the absence of any social targets on all days (including the conditioning days). A multifactor ANOVA consisting of two factors: zones (Inhibited, Control) and days (Baseline, Conditioning day 1, Conditioning day2, Test day) was used to determine significance.

Mice expressing NpHR/YFP in PL-NAc neurons, were also tested on a homecage assay. One of the two experimental days was designated as the inhibition day and the other as control day. On the inhibition day, mice were allowed to investigate a social target introduced into their homecage while PL-NAc neurons were inhibited with yellow light ( 594 nm, 2.5-3 mW). On the control day, mice explored investigated a novel social target while tethered but receiving no light stimulation. Half the mice received light on day 1 while the rest on day 2. A multifactor ANOVA, consisting of two factors: groups (ChR2, YFP) and days (Control day, Inhibition day) was used to determine significance.

### Endoscopic GCaMP6f imaging in freely moving mice

~3-4 weeks post injection, mice expressing GCaMP6f in PL-NAc neurons were acclimatized to being head-fixed and to carrying the microscope. A tethered dummy microscope was attached to the baseplate implanted on the animal for 30 min sessions across a 2-5 day period prior to the actual imaging experiment. On the day of testing, mice were left to habituate to the microscope for a 30 min window prior to imaging. Images were acquired at 10Hz and the LED power on the scope was set between 10-40% (0.3-2mW). A DAQ card (USB-201, Measurement computing) was used to synchronize the imaging and behavioral data.

For GCaMP6f imaging in the linear 3-chamber assay, all mice were imaged while exploring the arena on two consecutive visits (each visit being 10 min long): first with the social target in one location (either “social target right” or “social target left”), and next with the social target in the other location (e.g. Condition 1: social target on right, novel object on left; Condition 2: social targets mouse on left, novel object on right).

For imaging during the home cage assay, each imaging mouse was exposed to 2 male, novel conspecifics (“social target”) and a novel object presented in a random order in their home cage. GCaMP6f imaging and video monitoring preceded the introduction of the social target and the novel object by a minimum of 30s.

For data analysis, the imaging data was spatially downsampled (by a factor of 2 or 4) and motion corrected with a translational correction algorithm based on cross-correlations computed on consecutive frames using the Mosaic software (Inscopix, motion correction parameters: translation only, reference image: the mean image, speed/accuracy balance: 0.1, subtract spatial mean [r = 20 pixels], invert, and apply spatial mean [r = 5 pixels]). After motion correction, the CNMFe algorithm [77] was used to identify individual neurons, obtain their fluorescence traces and to deconvolve the fluorescence signal into firing rate estimates. These firing rates were used for spatial receptive field estimates in figures 4D-G, 5B-C and 6C-D. Custom MATLAB software was used for all imaging data analysis.

### Estimating spatial fields from Ca2+ imaging data

To estimate spatial fields and their modulation by social investigation (Figs 4D-G, 5B-C and 6C-D), we used a Bayesian approach based on a Gaussian process (GP) regression model [75] for calcium fluorescence imaging data. This model provides a principled statistical framework for quantifying the relationship between spatial position and neural activity.

There are several key advantages of this approach. First, it takes into account the slow dynamics of the calcium reporter so that neural activity will not be mis-attributed to the wrong location or action, as mice can travel a significant distance (or transition from social investigation to not investigation) during a single calcium transient. This also ensures that long calcium transients are not treated as providing independent samples of elevated firing, which will confound a naive shuffling-based permutation tests for significance of spatial tuning. Second, it automatically estimates the smoothness of spatial RFs using maximum likelihood, an approach known as “maximum marginal likelihood” [76]. This step is especially important for these data, given that we have no a priori knowledge about the smoothness of the spatial receptive fields of PL-NAc neurons. Third, taking a Bayesian approach allows us to quantify uncertainty about the influence of each behavioral variable on neural activity (in this case, social investigation and spatial position) in the form of a posterior distribution over model parameters. This is particularly valuable in the case of social investigation data, given that behavioral parameters of interest may co-vary, and each condition is not necessarily sampled equally thoroughly (e.g. social investigation vs not investigation), which could be an issue when combined with the low rate of gCaMP transients in PL-NAc neurons. However, it is worth noting that all results reported in the paper are considered significant based on cross-validation, further ensuring that spatial selectivity estimated using this approach is robust to model mis-specification (cross validation details below).

To fit the model to data with this Bayesian approach, we discretized the receptive field *f* into bins of size *X* (~3 cm), resulting in a discrete place field map with bins (42 bins in the linear 3-chamber assay and 247 bins in the home cage assay). The Gaussian process places a multivariate zero-mean Gaussian prior distribution over the receptive field, 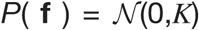, where the *i*, *j* ‘th element of the covariance matrix *K* is given by the “gaussian” covariance function: *K*(*i*,*j*) = *ϱ* exp (−||*x_i_* − *x_j_* ||^2^ / (2*δ*^2^)), where *x_i_* and *x_j_* are the spatial location of the *i*’th and *j*’th bins of the receptive field, respectively. This prior is governed by a pair of hyperparameters: marginal variance *ϱ*, which defines the marginal prior variance of the receptive field elements, and length scale *δ*, which determines the degree of smoothness. We assume neural activity in each time bin given the animal’s location follows a Gaussian distribution, 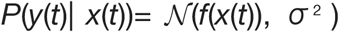, where *y*(*t*) is the neural activity at time *t*, *x*(*t*) is the animal’s spatial location at time *t*, and the distribution over activity has mean *f*(*x*(*t*)) and variance *σ*^2^. We set the Gaussian process hyperparameters *ϱ* and *δ* and Gaussian noise variance *σ*^2^ by maximizing marginal likelihood, the conditional probability of the data given these hyperparameters. This allowed us to determine the optimal degree of smoothing for each receptive field directly from the data. Then, conditioned on the hyperparameters, we used the maximum a posteriori (MAP) estimate to determine the receptive field for each neuron using standard formulas for the posterior under Gaussian prior and likelihood: f = (*X^T^ X* + *K*^−1^)^-1^ *X^T^ Y* where *X* is the design matrix, in which each row carries a 1 in the column corresponding to the spatial location of the rat in a single time bin, and the corresponding element of the column vector *Y* is the neural response for that time bin. This Matlab code will be uploaded to github upon publication.

To assess the significance of each neuron’s spatial selectivity in the linear 3-chamber assay (Figs 4D-G, 5B-C), a cross-validation method (based on leaving out 10% of the data) was used to compare the predicted fluorescence (based solely on the spatial receptive field) with the actual fluorescence. Neurons were classified as having ‘significant’ responses when there was significant correlation between the predicted and actual fluorescence (p<0.05).

For the data in Figure 4G, to determine if there were more “spatial neurons” and “social neurons” than expected by chance, the distribution of the peak activity was compared to what would be expected if peak responses followed a spatially uniform distribution. The number of neurons (from the real data set) in each bin was compared to simulated ata using a1-tailed t-test and bins with p values < 0.05 were designated as significant.

For the data in Figure 6C-D, the imaging and behavioral data were concatenated across exposure of two social target mice. To determine if adding social investigation as a parameter improved our prediction of the activity of each neuron, we compared two models. One model was based on neurons having the same spatial receptive field regardless of whether or not the mice was interacting (spatial-only model), and the other model was based on calculating a different spatial receptive field during investigation versus during non-investigation epochs (social+spatial model; smoothness parameters were determined from the entire dataset). Cross-validation (based on leaving out 1% of the data) was used to determine how the predicted fluorescence based on each model compared to the actual fluorescence. Correlation coefficients comparing the predicted and actual fluorescence were generated for each neuron and for both models. A paired t-test was then used to compare the correlation coefficients generated by the two models. Neurons were classified as “social+spatial” or “spatial” in Fig. 6E based on which model led to the greatest correlation coefficient with cross-validation.

## FIGURES

**Supplementary figure 1.**
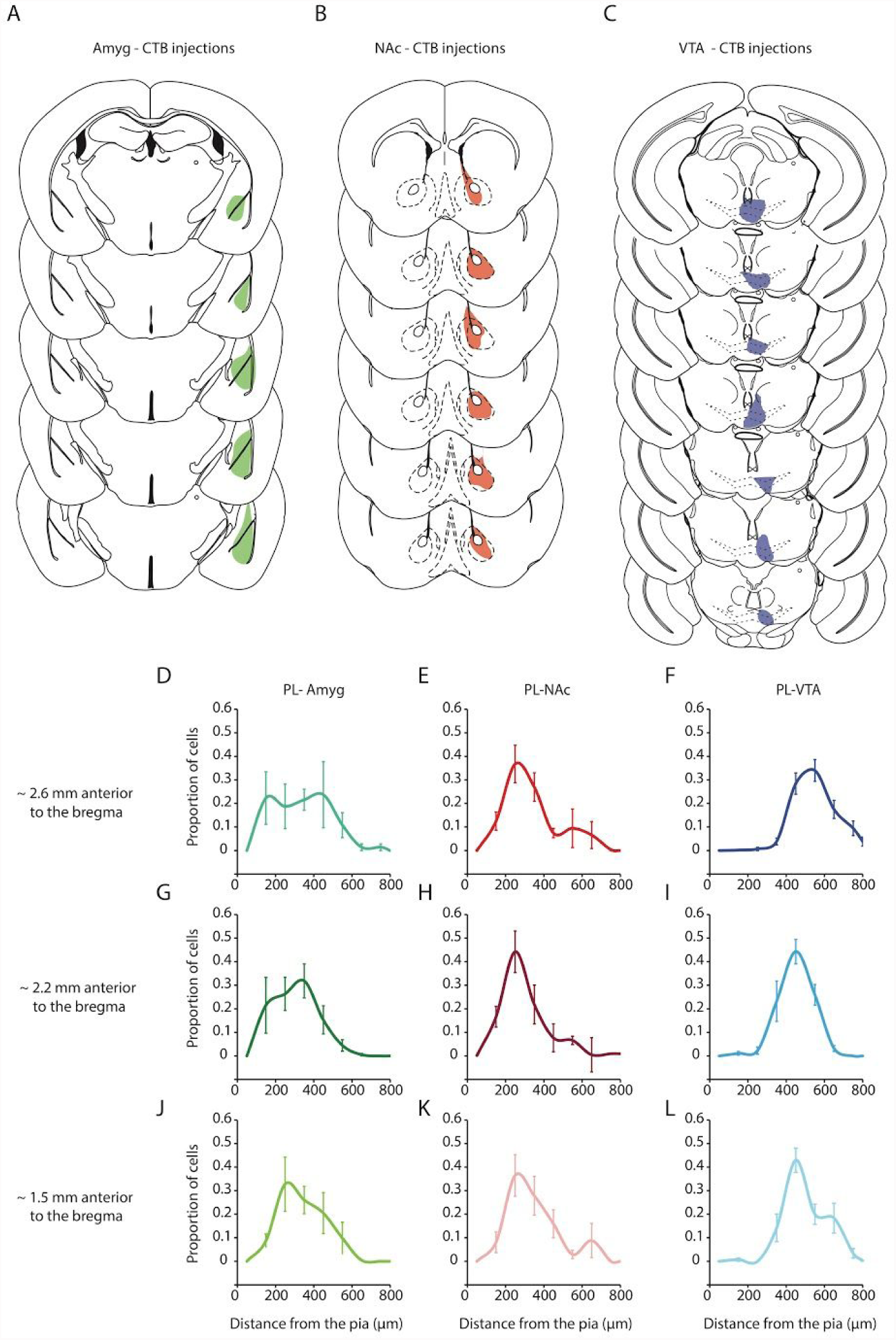
PL neurons projecting to the NAc, VTA and Amyg occupy distinct cell layers along the anterior-posterior axis. **A-C.** Reconstructions of injection sites (determined by maximal spread) in mice injected with CTB in the Amyg (A; green), NAc (B; red) and the VTA (C; blue). **D.** Distribution of PL-Amyg (left column, shades of green), PL::NAc (middle column, shades of red) and PL::VTA (right column, shades of blue) projection neurons along the anterior-posterior axis (top row: ~2.6mm anterior to bregma, middle row: ~2.2mm anterior to bregma and bottom row: ~1.5 mm anterior to the bregma). Error bars denote S.E.M.

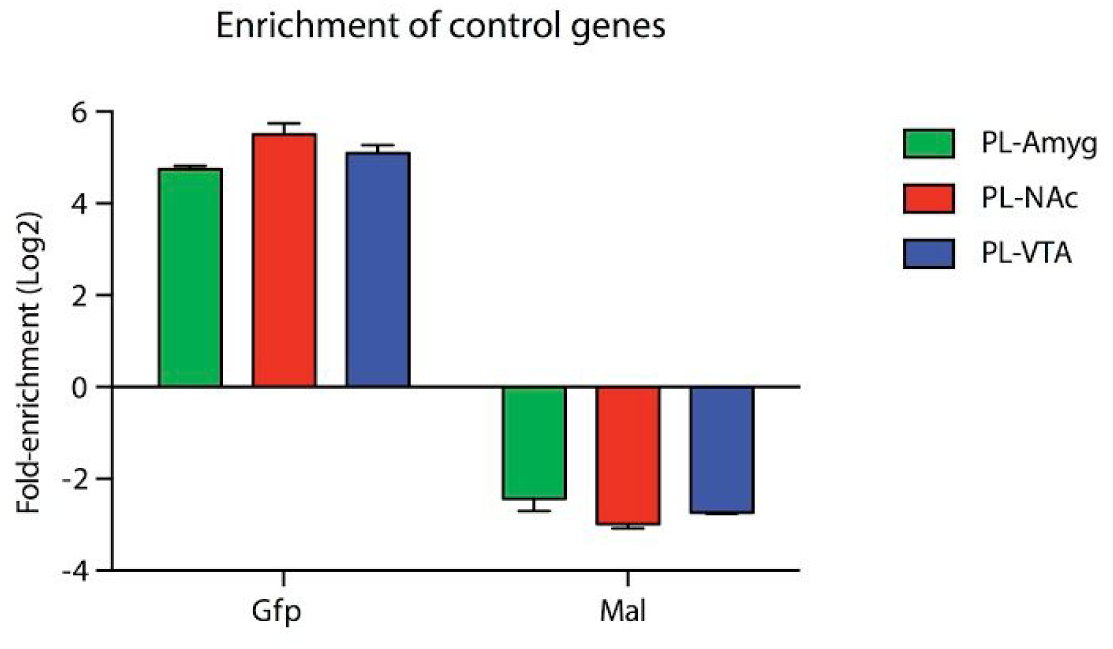
Supplementary figure 2. Enrichment of control marker genes across the 3 PL subpopulations. Enrichment of the positive control marker Gfp and a depletion of the glial cell marker Mal was observed across all three PL subpopulations. Fold enrichments (IP/Input) are on a Log2 scale.

**Supplementary figure 3.**
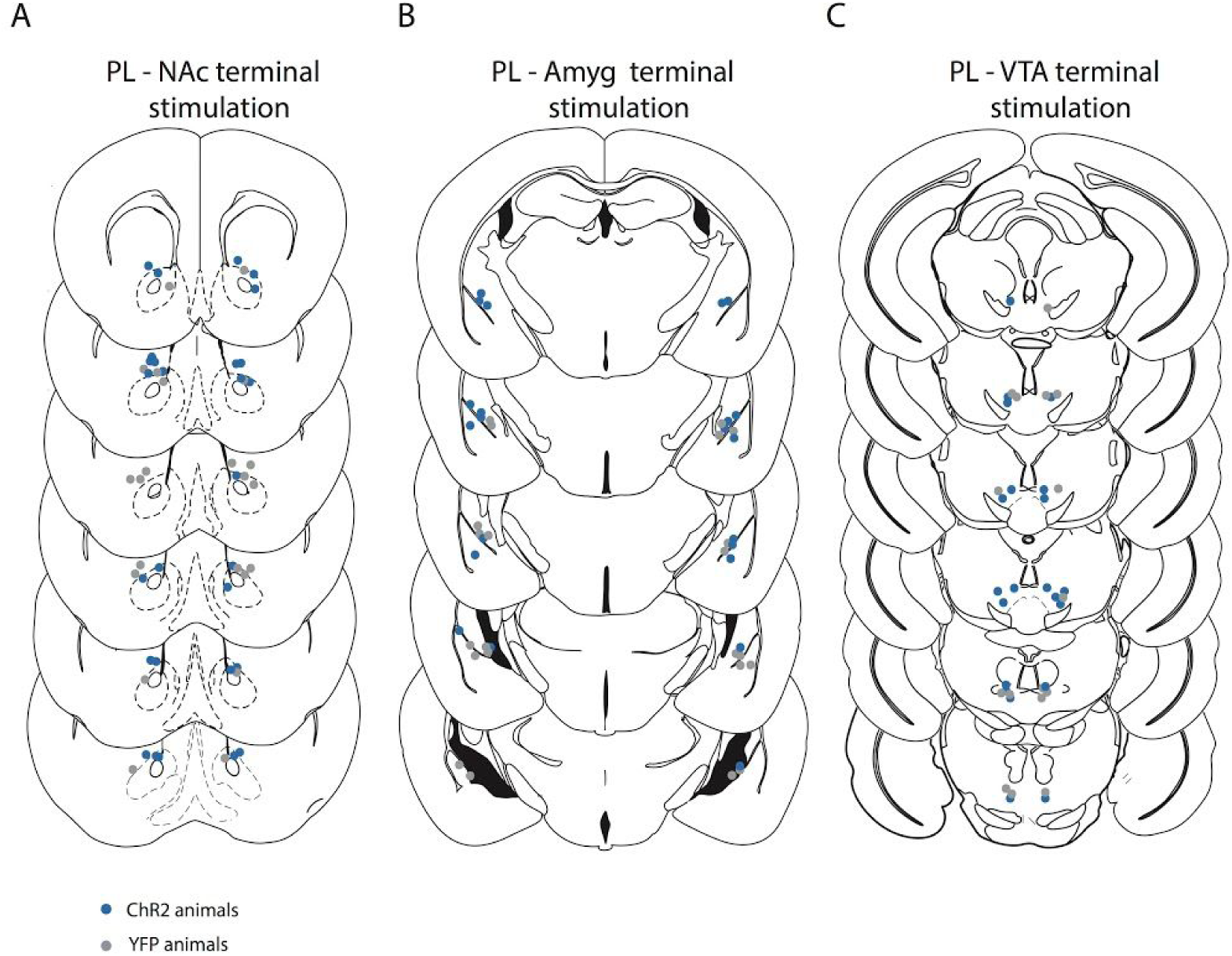
Location of optical fiber tips targeting PL terminals in the NAc, Amyg and VTA. **A.** Histological reconstruction of optical fiber tip placements in the PL-NAc mice, (blue dots: ChR2, grey dots: YFP). **B.** Histological reconstruction of optical fiber tip placements in the PL-Amyg mice. **C.** Histological reconstruction of optical fiber tip placements in the PL-VTA mice.

**Supplementary figure 4.**
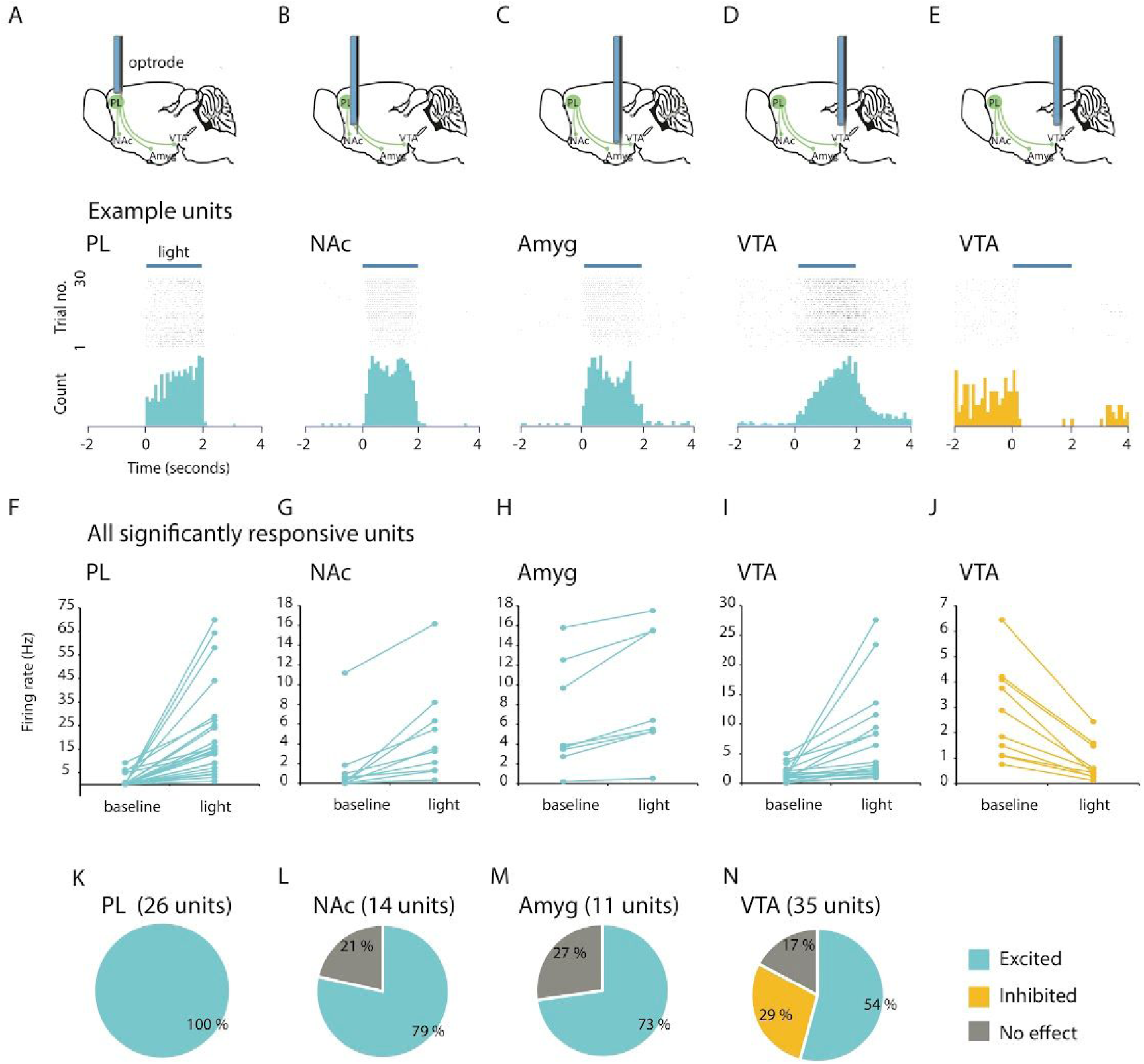
Electrophysiological confirmation of ChR2-mediated activation of PL terminals in the NAc, Amyg and VTA. **A.** *In vivo* extracellular recording of light-evoked action potentials (473nm, 2s, 30 trials) in the PL of mice injected with AAV5-CamKIIa-hChR2-EYFP in PL. Top row: schematic of the recording configuration, optrode inserted in the PL. Middle row: Raster plot showing action potentials of an example unit in PL in responding to 30 consecutive trials of light illumination. Bottom row: A histogram of the spikes binned every 100 ms. **B.** Example of a NAc unit excited by the activation of PL terminals. (Top row: Schematic, Middle row: Raster plot, Bottom row: Histogram). **C.** Example of an Amyg unit excited by the activation of PL terminals. **D.** Example of a VTA unit excited by the activation of PL terminals. **E.** Example of a VTA unit inhibited by the activation of PL terminals. **F.** Population summary of firing rate changes of all PL units recorded that showed significant increases from the baseline (P<0.0001, Paired two tailed t-test comparing mean firing rate across epochs). **G-I.** Population summary of firing rate changes of units in the NAc, Amyg and VTA that showed significant increases in their firing rate during stimulation of ChR2 expressing PL terminals (NAc: p=0.003, Amyg: p=0.004, VTA: p=0.005, for Paired two tailed t-test). **J.** Population summary of firing rate changes of units in the VTA that showed significant decreases in their firing rate during stimulation of ChR2 expressing PL terminals (p=0.006, Paired two tailed t-test). **K-N.** Percentage of PL (**K**), NAc (**L**), Amyg (**M**) and VTA **(N**) units excited (blue), inhibited (yellow) or not affected (grey) by light stimulation.

**Supplementary figure 5.**
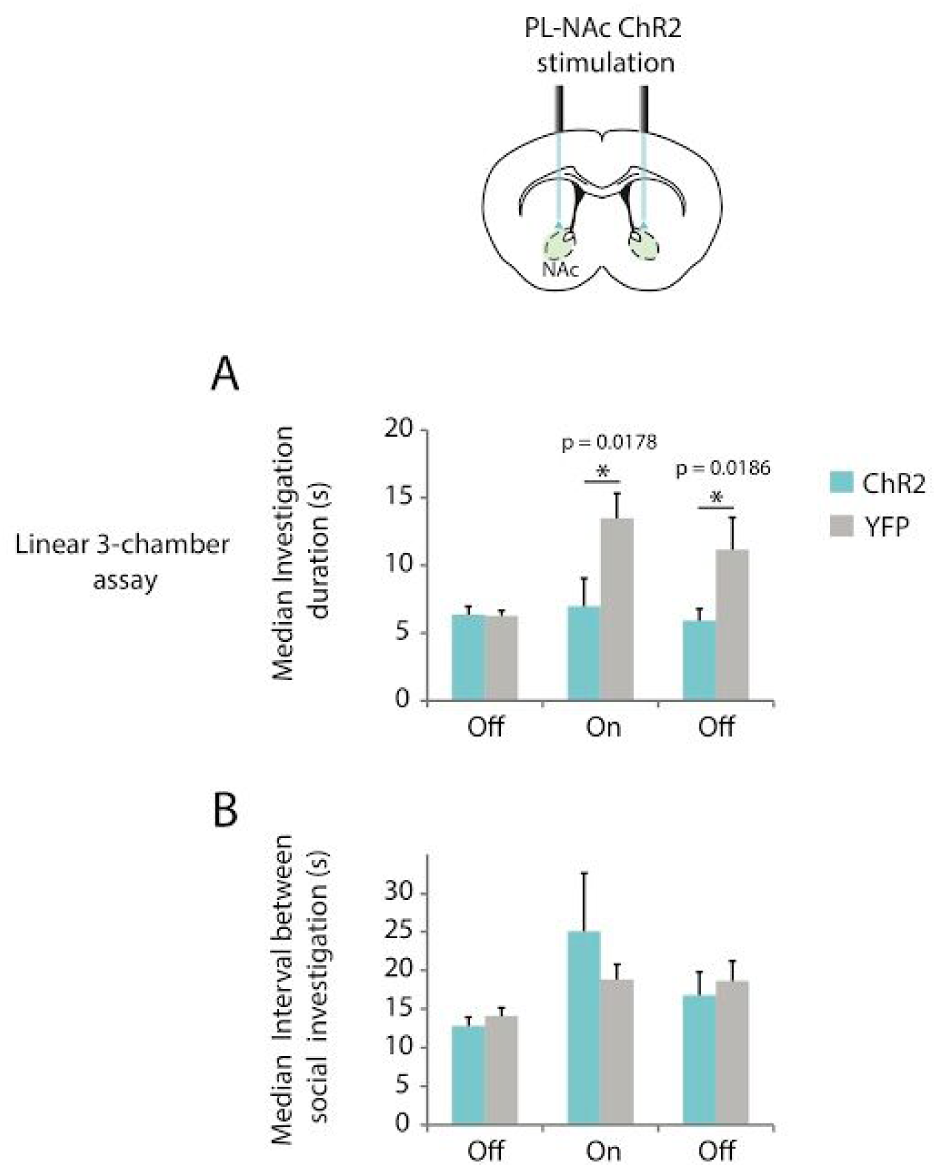
Activation of PL-NAc terminals expressing ChR2 disrupts social investigation. **A.** Stimulation of PL-NAc terminals decreased the duration of social investigation bouts and this effect persists after light was turned off. (ANOVA with group, epoch, and light as factors; *p* = 0.049, ANOVA group x light interaction). **B.** The stimulation of PL-NAc terminals had no effect on the intervals between social investigation bouts (*p*= 0.505, ANOVA group x light interaction).

**Supplementary figure 6.**
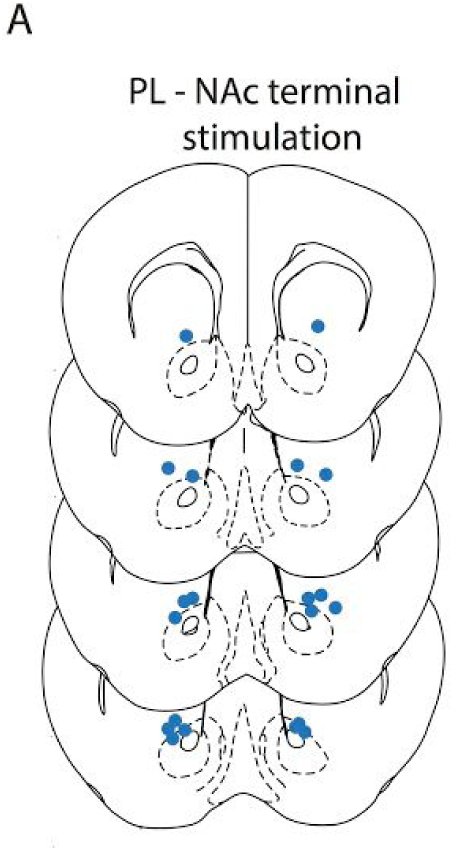
Location of optical fiber tips targeting PL terminals in the NAc from mice used in the home cage assay. **A.** Histological reconstruction of optical fiber tip placements in the PL-NAc mice, (blue dots: ChR2, n=10 animals)

**Supplementary figure 7.**
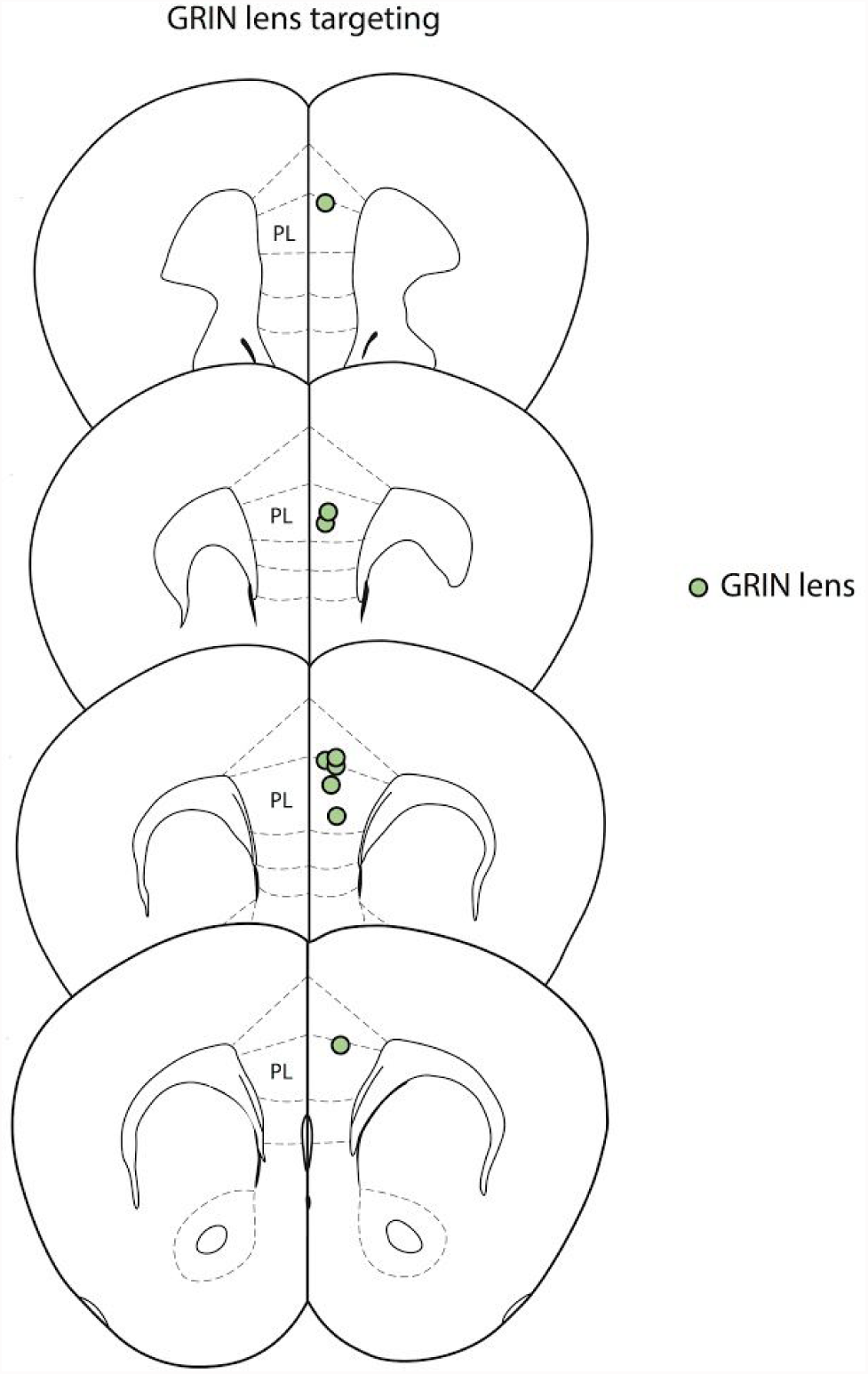
Reconstructing GRIN lens tip locations targeting PL-NAc neurons. Histological reconstruction of GRIN lens tip placements in the PL of mice expressing GCaMP6f in PL-NAc projection neurons (green dots, n=9 animals).

**Supplementary figure 8.**
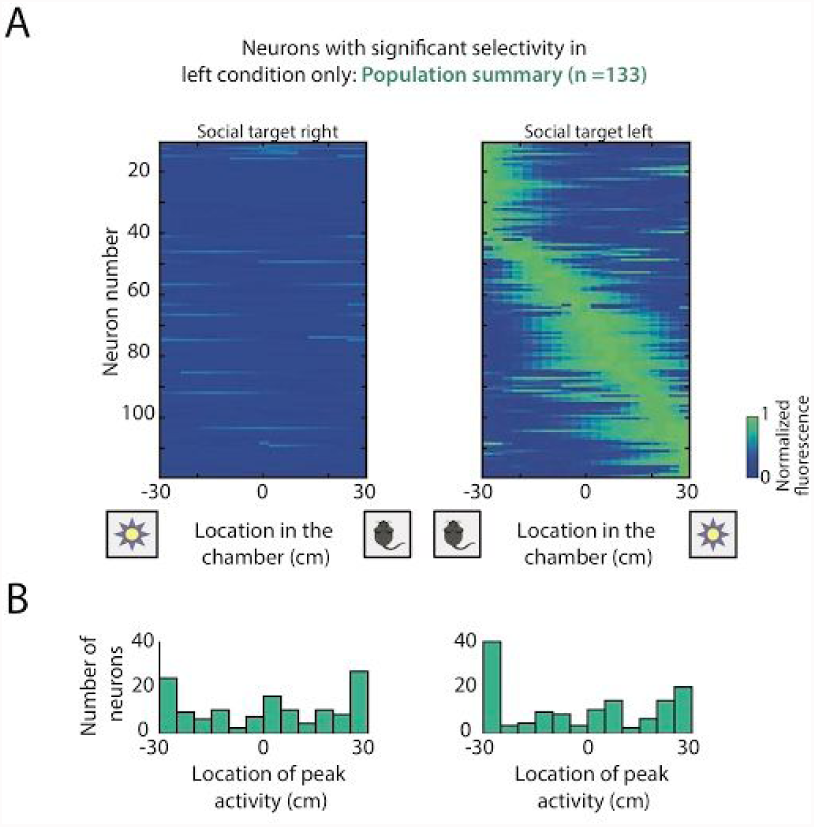
In a 3-chamber assay, many PL-NAc neurons respond in the vicinity of the social target, only in a specific spatial location. **A.** Spatial receptive fields of all neurons with significant responses (n=133) only in the social target left condition (right panel) and not the social target right condition (left panel). **B.** Histogram of the peak responses of neurons along the length of the chamber showing many neurons responding in the proximity of the social target in the social target left condition (right panel), while no distinction between the social target and novel object was observed in the non-preferred social target right condition (left panel).

